# Constrained release of lamina-associated enhancers and genes from the nuclear envelope during T-cell activation facilitates their association in chromosome compartments

**DOI:** 10.1101/062224

**Authors:** Michael I. Robson, Jose I. de las Heras, Rafal Czapiewski, Aishwarya Sivakumar, Alastair R. W. Kerr, Eric C. Schirmer

**Affiliations:** The Wellcome Trust Centre for Cell Biology and Institute of Cell Biology, University of Edinburgh, Edinburgh EH9 3BF, UK

**Author notes:** Correspondence to: Eric Schirmer Wellcome Trust Centre for Cell Biology University of Edinburgh, Kings Buildings Swann 5.22, Max Born Crescent Edinburgh EH9 3BF, UK.

## Abstract

**Abstract:** The 3D organization of the genome changes concomitantly with expression changes during hematopoiesis and immune activation. However, studies of this phenomenon have focused either on lamina-associated domains (LADs) or on topologically-associated domains (TADs), defined by preferential local chromatin interactions, and chromosome compartments, defined as higher-order interactions between TADs in functionally similar states. However, few studies have investigated how these affect one another. To address this, we mapped LADs using Lamin B1-DamID during Jurkat T-cell activation, finding significant genome re-organization at the nuclear periphery dominated by release of loci frequently important for T-cell function. To assess how these changes at the nuclear periphery influence wider genome organization, our DamID datasets were contrasted with TAD contact domains and compartments identified with high resolution in the GM12878 lymphoblastoid cell line. Features of specific repositioning events were then tested by fluorescence *in situ* hybridization. First, considerable overlap between TAD contact domains and LADs was observed and we found that the TAD repositioned as a unit during T-cell activation. Second, sub-compartments of the A compartment, termed A1 and A2, are segregated in 3D space through differences in proximity to LADs along chromosomes. Third, genes and a putative enhancer that occurred in LADs that were released from the periphery during T-cell activation tended to become associated particularly with A2 sub-compartments and were thus constrained to the relative proximity of the lamina. Thus, lamina-associations clearly influence internal nuclear organization while changes in LADs during T-cell activation may provide an important additional mode of gene regulation.

## Introduction

The genome is non-randomly organized within the three-dimensional space of the interphase nucleus. At one level, regulatory elements influence the activity of target genes many hundreds of kbp to many Mbp distal by moving into close proximity through looping in three-dimensional space (Sanyal et al. 2012; Shen et al. 2012; Arner et al. 2015; Schoenfelder et al. 2015). Chromosomes are also organized along their length into discrete large genomic regions termed topologically associated domains (TADs) that display more frequent interactions based on chromosome conformation capture (Hi-C) approaches within the domain than with the rest of the genome (Dixon et al. 2012; Hou et al. 2012; Nora et al. 2012; Sexton et al. 2012). In general, the topological constraints imposed by TADs limit the propensity of looping interactions to *cis*-regulatory element:gene pairings present within the same domain (Lupianez et al. 2015; Franke et al. 2016; Hnisz et al. 2016). However, a higher level of TAD organization also occurs in which multiple TADs sharing a common functional state that are many Mbp distal on the same chromosome or on different chromosomes can also be adjacent in the higher order folding of the genome (Lieberman-Aiden et al. 2009). These associating TADs are referred to as compartments, which segregate into two classes; A, correlating with active transcription, and B, correlating with a repressive state (Lieberman-Aiden et al. 2009). While the composition of TADs is largely invariant, the composition of compartments has been shown to change during tissue differentiation and between different cell types in a manner correlated with gene activity (Dixon et al. 2015; Fraser et al. 2015); however, how interactions between compartments on the same chromosome are controlled during differentiation or other cellular transitions is largely unexplored.

A completely different level of radial genome organization occurs with respect to the nuclear periphery. The proximity of DNA regions with the nuclear lamina, a proteinaceous meshwork of polymers of nuclear lamin intermediate filaments that lines the inner surface of the nuclear membrane, generates so-called lamina-associated domains (LADs) that define genome regions at the nuclear periphery (Guelen et al. 2008). However, while providing physical support for the nucleus and being implicated in genome organization (Mewborn et al. 2010; Mattout et al. 2011; Solovei et al. 2013), lamins themselves are not absolutely required for tethering LADs to the periphery (Amendola and van Steensel 2015). The nuclear periphery is a generally repressive environment with genes in LADs tending to overlap with late-replicating DNA lacking active histone marks and containing many silencing marks (Guelen et al. 2008). Many LADs are constitutive, occurring in all cells investigated thus far, but as more cell types are investigated the subset of facultative LADs that change in their peripheral associations either during the cell cycle or senescence or between cell types is constantly increasing (Peric-Hupkes et al. 2010; Kind et al. 2013; Meuleman et al. 2013; Chandra et al. 2015; Robson et al. 2016). The importance of facultative LADs is underscored by the fact that knockdown of nuclear membrane proteins involved in the establishment of some facultative LADs has profound negative consequences for tissue differentiation (Robson et al. 2016). Tellingly, these facultative LADs often contained genes requiring fine-tuned regulation in differentiation and involved genomic regions both moving to and from the periphery during differentiation (Peric-Hupkes et al. 2010; Robson et al. 2016).

Though LADs have been shown to correlate with TADs and specifically repressed compartments (Dixon et al. 2012; Rao et al. 2014; Fraser et al. 2015), few studies have investigated how alterations to one can influence the others. Several questions come to mind. Does a TAD as a functional unit move between the periphery and interior and thus perhaps encompass or be contained within a facultative LAD? Can a new LAD that forms during differentiation influence the organization of TADs or compartments or vice-versa?

The movement of genes in facultative LADs between the nuclear periphery and the nuclear interior tends to be associated with their repression or activation respectively. In neurogenesis and myogenesis 5-15% of genes move from the nuclear periphery to the interior while another 5-10% move from the interior to the periphery (Peric-Hupkes et al. 2010; Robson et al. 2016). Though similar DamID datasets have not been presented for lymphocyte differentiation, several specific gene-positioning changes have been observed by fluorescence in situ hybridization (FISH) such as release of the *IgH* locus from the periphery upon induction of *V-D-J* recombination (Kosak et al. 2002) and movement of the *c-maf* locus to the periphery with its repression (Hewitt et al. 2004).

Lymphocyte activation in contrast to differentiation is a much more rapid and dynamic process associated with massive genome restructuring (Drings and Sonnemann 1974). Compacted chromatin in the resting stage dissipates in the activated state concomitant with large-scale gene activation (Pompidou et al. 1984). This raised the question of whether the massive gene activation and chromatin decondensation in lymphocyte activation is whole-scale and unidirectional or whether there is a specifically regulated exchange of genes tethered at the nuclear envelope as in differentiation. We anticipated that some level of regulation occurs from the nuclear envelope because changes in the protein composition of the nuclear membrane were observed during lymphocyte activation (Korfali et al. 2010). To address these questions we have globally identified changes in genome contacts with the nuclear periphery in resting and activated Jurkat T-cells and compared the genes changing position with a published high-resolution Hi-C dataset, confirming several hypotheses raised by this analysis using FISH. We find that indeed there is a specific exchange of genome contacts as opposed to a whole-scale release from the periphery and that upon lymphocyte activation specifically released and induced genes and a predicted enhancer can associate in a compartment constrained near the nuclear periphery.

## Results

### Mapping gene expression and repositioning changes during T-cell activation

The extensive electron dense peripheral heterochromatin found in resting T-cells dissipates during activation concomitantly with the induction of immunogenic genes (Hirschhorn et al. 1971; Pompidou et al. 1984; Manteifel et al. 1992; Rawlings et al. 2011). However, it is unclear whether amongst what appears to be a whole-scale reorganization of the genome there is more specific functional regulation for immune activation. We predicted that the dissipation of peripheral heterochromatin would specifically correlate with a release of T-cell activation-associated genes from the periphery. Accordingly, we investigated coordinated gene expression and genome organization changes during T-cell activation using microarrays and DamID (Fig. 1). Jurkat cells were incubated with Raji B-cells that had been pre-conjugated with staphylococcal enterotoxin E (SEE) resulting in the formation of antigen-independent immunological synapses as has been reported previously (Fig. 1A) (Gonzalez-Granado et al. 2014). This yielded robust activation of over 95% of Jurkat cells as assessed by CD69/CMAC fluorescence activated cell sorting (FACS) (Fig. 1B). To determine gene expression changes associated with activation, RNA was extracted at 0, 8, 24 and 48 h post-SEE stimulation and analyzed on Illumina bead microarrays. Upon activation, 1,111 genes were upregulated at least 1.4-fold at any of the three time points and these were significantly enriched in Gene Ontology (GO)-terms positively supporting T-cell activation and early effector function (Figure 1C, Supplemental Fig. S1). Similarly, 1,016 genes were repressed at least 1.4-fold and these were significantly enriched in GO-terms inhibiting mitosis and cell division, presumably in order to permit accelerated proliferation of activated cells. Thus, both FACS and GO-term analysis confirmed efficient Jurkat T-cell activation.

**Figure 1.**
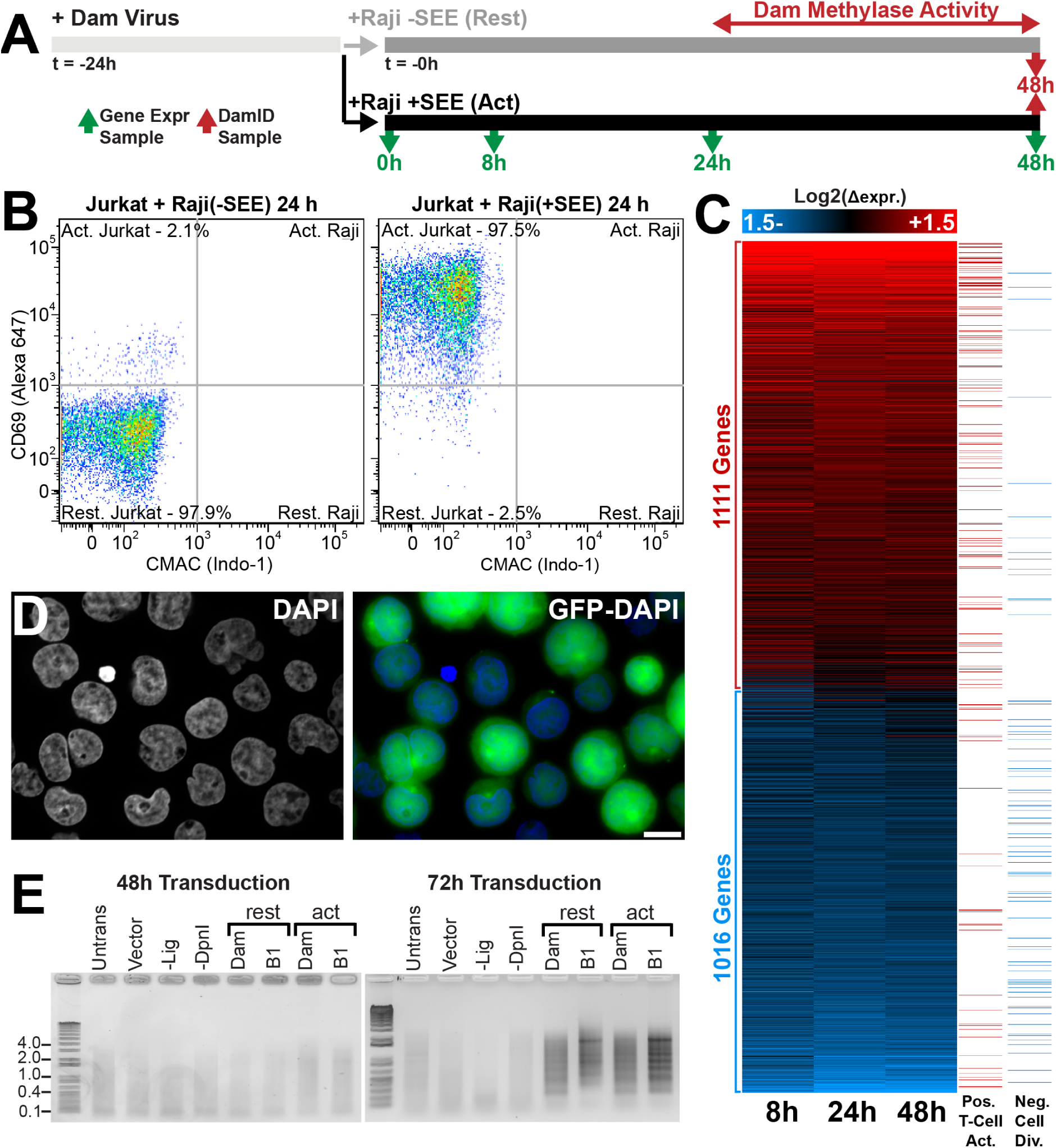
Determination of gene expression and genome organizational changes during T-cell activation. (*A*) Schematic diagram demonstrating methodology for microarray gene expression and DamID analysis of Resting and Activated Jurkat T-cells. (*B*) FACS plot of CD69 vs CMAC staining for Resting Jurkat T-cells either mock-activated with Raji B-cells lacking antigen or activated with Raji B-cells presenting SEE antigen. The Raji B-cells were gated out after staining with CMAC to distinguish the antigen-presenting cell from Jurkat T-cells. 97.5,% of Jurkat T-cells become activated when stimulated with SEE conjugated Raji B-cells. (*C*) Heat map of gene expression changes of at least 1.4 fold for the three timepoints taken with GO terms associated with positive regulation of T-cell activation or negative regulation of cell division highlighted in the right-most panels. See also Supplemental Fig. S1 for associated GO-terms. (*D*) Representative micrograph of Jurkat T-cells transduced with a GFP-encoding lentivirus indicating a likely high transduction efficiency for the parallel transduced DamID constructs. (*E*) Agarose gels of PCR amplified Dam methylated genomic DNA. Amplification is only observed in Resting or Activated cells 72 h post transduction. T4 DNA ligase null (-Lig) and *DpnI* null (*-DpnI*) controls represent background amplification. For DamID analysis results are derived from a single experiment while gene expression analysis was performed in triplicate.

The same cell populations were subjected to lamin B1-DamID to map global changes in associations between genomic loci and the nuclear periphery in resting and activated Jurkat cells. Bacterial Deoxyadenosine methylase (Dam) fused to lamin B1 was delivered to cells by lentiviral transduction where it could provide its unique methylation to DNA proximal to the lamin polymer that underlies the inner nuclear membrane (Vogel et al. 2007). The methylated DNA was then isolated and identified by sequencing. To control for local variation in chromatin accessibility, in parallel experiments Dam methylase expressed independently of lamin B1 so that it distributed throughout the nucleoplasm (Dam-only) was analyzed. Although a GFP reporter lentivirus delivered in a parallel sample confirmed efficient transduction (Fig. 1D), time course experiments revealed no DamID signal until roughly 48 h after lentiviral transduction (Fig. 1E). Hence, Jurkat cells were transduced with DamID constructs 24 h before antigen presentation so that the measured changes reflect the genome organization between 24 and 48 h into Jurkat activation (Fig. 1A).

Log2(Lamin B1 Dam/Dam-only) ratios were generated and used to identify lamina associated domains (LADs) in resting and activated Jurkat T-cells based on a circular binary segmentation algorithm with slight modifications from that previously described ((Harr et al. 2015); see materials and methods). The LADs were mapped onto the hg19 assembly because other datasets used for comparison were also mapped on this assembly, but all regions examined by FISH (see below) have been confirmed to be unaltered between the hg19 used here and the more recent hg38 assembly. Interestingly the magnitude of changes were similar between this rapid activation system and previous DamID studies of differentiation investigating neurogenesis and myogenesis (Peric-Hupkes et al. 2010; Robson et al. 2016) with the majority of LADs maintained between resting and activated cells. 96% of the LAD coverage was shared between resting and activated cells with the remaining 4% roughly equally distributed between lost and newly formed LADs (summarized in Figures 2A and 3A). Moreover, the 18.6% of genes found within LADs were less transcriptionally active than the 81.3% of genes found in non-LADs (Fig. 2B), in line with other studies representing the periphery as a repressive environment (Pickersgill et al. 2006; Pindyurin et al. 2007). Similar differences in gene expression between LAD and non-LAD compartments were also observed when the Jurkat DamID data was compared with expression data from primary cells taken from a published naïve CD4^+^ RNA-seq dataset (Kundaje et al, 2015; Supplemental Fig. S2A). Hence, although being an immortalized T-cell acute lymphoblastic leukemia cell line, Jurkat T-cells and naïve CD4^+^ T-cells display a broad correspondence between their gene expression state and lamina-association.

**Figure 2.**
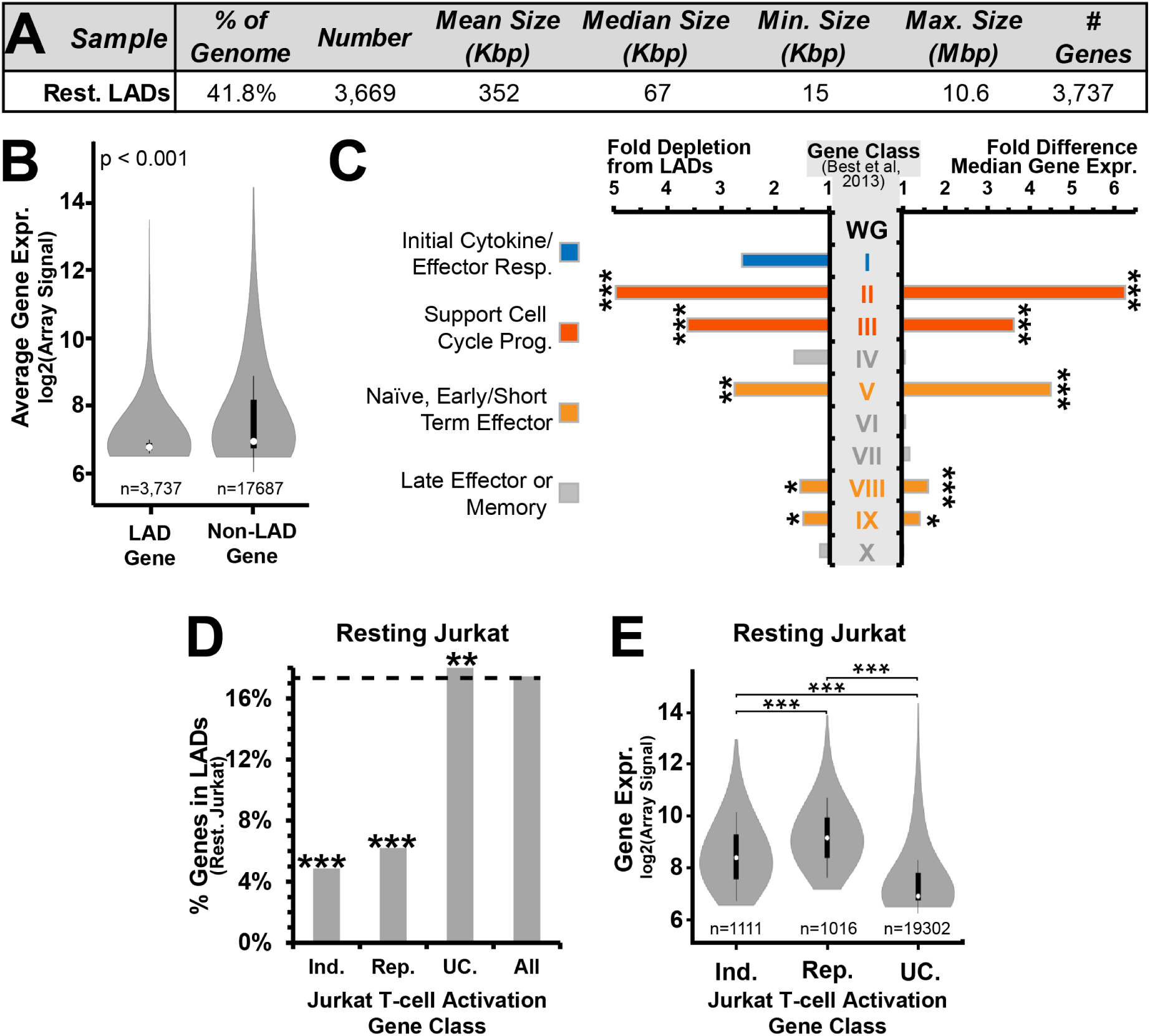
The frequency of lamina association is consistently related to gene expression and function. (*A*) Table summarizing parameters of identified resting LADs. (*B*) Violin plot showing the microarray determined gene expression level of genes within LADs compared to genes within non-LAD regions. (*C*) Fold differences in gene expression and frequency of lamina-association in resting Jurkat T-cells for 10 published categories of genes displaying distinct transcriptional behaviors during T-cell activation (Best et al. 2013). Classes associated with naïve cells, early effector function and cell division are more expressed and observed less frequently in LADs than classes associated with late effector/memory function. Further details of classes can be found in Figure S2. Fold differences were calculated relative to corresponding whole genome (WG) measurements in resting Jurkat T-cells. Higher expressed genes tend to be depleted from LADs. (*D*) Bar plot showing the percentage of genes occurring in LADs in resting Jurkat T-cells for different expression categories: genes induced (Ind.) or repressed (Rep.) at least 1.4 fold during T-cell activation, unchanged genes (UC.) and all genes in resting Jurkat T-cells. Both induced and repressed genes are 3-4 times less frequently found in LADs compared to genes that are not altered in expression. (*E*) Violin plot for the expression level distribution of induced, repressed, or unchanging genes. Both induced and repressed categories are more highly expressed than the unchanging genes in resting Jurkat T-cells. For B, C expression analysis and E, significance of difference in microarray gene expression was determined by Dunn test for multiple comparison testing after a Kruskal Wallis significance test. For C and D frequency of lamina-association analysis, statistical significance was determined by Fisher tests. \**P* < 0.05, \*\**P* < 0.01 and \*\*\**P* < 0.001. See also Supplemental Fig. S2 and S3 and Supplemental Table S1.

### Gene expression and function reflects the frequency of lamina-association

On a genome-wide scale the frequency of lamina-association relates to the level of gene expression (Fig. 2B). However, as LADs are enriched in developmentally regulated loci (Peric-Hupkes et al. 2010), we next sought to determine if this genome-wide relationship applied to individual functional classes of genes. To assess this, we focused on a selection of 10 different classes of genes (Supplemental Fig. S2B) that each displayed distinct transcriptional dynamics in an extensive published time course of mouse CD8^+^ T-cell activation (Best et al. 2013). The frequency of lamina-association in resting Jurkat T-cells was compared with the expression of the genes from these classes in resting Jurkat T-cells (Fig. 2C). This revealed that 5 gene classes involved in early stages of T-cell activation, such as cell cycle progression and early/short term effector function, were significantly depleted from LADs relative to the whole genome and displayed increased gene expression. In contrast, 4 gene classes not needed until later in T-cell responses, such as those associated with late effector and memory T-cell function, were found in LADs as frequently as the whole genome and were mostly transcriptionally repressed whether in LADs or not. These relationships are largely maintained when instead considering expression data from naïve CD4^+^ T-cells (Supplemental Fig.S2C). However, of the 5 gene classes depleted from LADs in resting Jurkat cells, two classes associated with active cell division and short term effector functions failed to display similarly elevated gene expression in naïve CD4^+^ T-cells (Supplemental Fig. S2C). Also, one class associated with naïve memory precursor function displayed elevated transcriptional activity in the naïve CD4^+^ T-cells, which was absent in resting Jurkat T-cells. The differences likely reflect properties of the Jurkat T-cells as a cancer cell line, for example the reason for active cell division gene classes being depleted from LADs in the resting Jurkat cells likely reflects their active cell division compared to a strong suppression of cell division in primary resting human naïve T-cells. Accordingly, these LAD-depleted gene classes were also expressed in the resting Jurkats.

To investigate the correspondence between lamina-association and gene expression further, we performed a similar analysis on all genes displaying induction and repression in Jurkat T-cell activation. As expected, genes that are repressed during lymphocyte activation are active in the resting cells and exhibit depleted lamina-association in resting cells, but surprisingly we also observed that induced genes were similarly excluded from LADs (Fig. 2D). However, although displaying elevated expression upon Jurkat T-cell activation, the induced genes also displayed elevated basal expression in the resting state relative to the whole genome (Fig. 2E) and this was recapitulated in the naïve CD4^+^ T-cell data (Supplemental Fig. S2D). This basal expression explains why they are depleted from LADs. These trends are similar to changes in myogenic differentiation as plotting data from a previous DamID study (Robson et al. 2016) revealed depleted lamina-association and elevated gene expression in undifferentiated myoblasts for genes normally induced or repressed during myogenesis (Supplemental Fig. S3A and B). Hence, depletion in lamina-association invariably corresponds to increased gene expression for all gene classes tested and in both lymphocyte activation and myogenic differentiation.

### Important T-cell genes are released from the periphery and induced during T-cell activation

Having characterized LADs in resting cells, we next sought to determine how gain or loss of lamina-association correlates with gene expression changes during T-cell activation. To ensure repositioning events to or from the periphery were identified with high confidence, we employed a stringent binary LAD or non-LAD definition. Whereas ∼ 40% of the genome could be found in LADs in both resting and activated Jurkat T-cells, only ∼2-2.5% of the genome was found uniquely associated with the lamina in only one of the conditions (Fig. 3A). A tendency to increase association with the periphery in activated cells versus resting cells (denoted interior to periphery ‘IP regions’) was observed for 1.9% of the genome in 2,251 regions with a median size of 22 kbp containing 554 genes. The opposite tendency (decreasing association from the periphery; denoted periphery to interior ‘PI regions’) was observed for 2.5% of the genome in 2,753 regions with a median size of 23 kbp, containing 792 genes (Fig. 3A). In addition to these parameters, minimum and maximum sizes of all LADs and differential IP/PI LAD regions were also calculated and summarized in Figure 3A. An example of the log2(B1/Dam-only) signal intensity shift and corresponding PI regions is shown in Figure 3B. Consistent with the dissipation of peripheral heterochromatin, more of the genome displays decreased association with the transcriptionally repressive nuclear periphery than increased association during T-cell activation.

**Figure 3.**
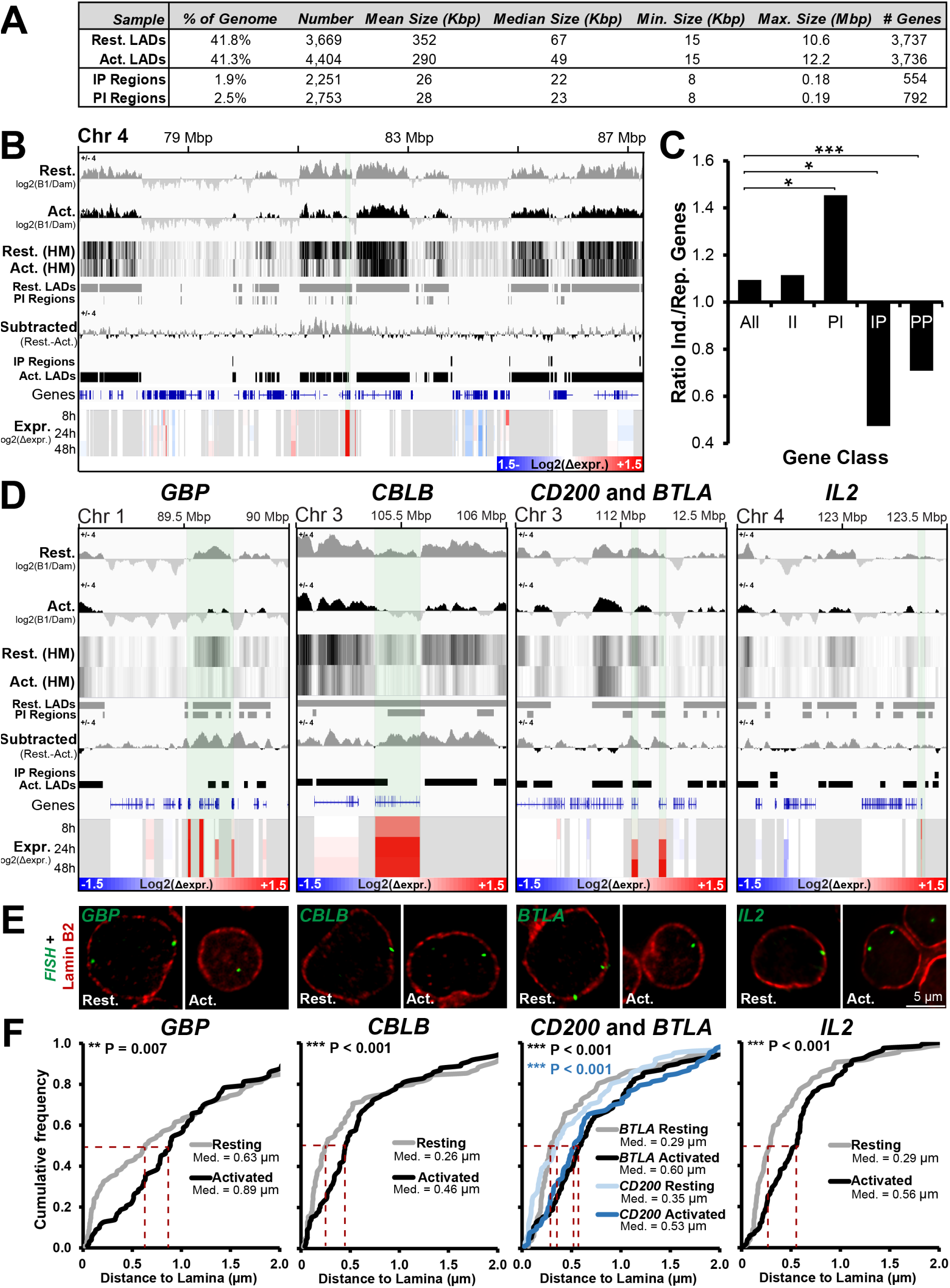
A number of important genes are released from the periphery during T-cell activation. (*A*) Table summarizing parameters of identified LADs and regions showing increased (IP) and decreased (PI) association with the nuclear periphery between resting and activated Jurkat T-cells. (*B*) Genome browser view for the genomic region surrounding *ANTRX2* showing DamID signal intensities as both bar plot and heat map (HM), identified LADs, IP and PI regions and microarray gene expression changes in Jurkat T-cells during activation from the resting state at 8, 24, and 48 h. The key for intensity of log2(Δexpr.) is given in the bottom right corner of the expression data. For clarity, the Resting-Activated subtracted DamID signal is also shown and the ANTRX2 gene position is noted by shading in light green. (*C*) Bar plot demonstrating ratio of genes induced (Ind.) and repressed (Rep.) during T-cell activation for the whole genome (All), PI, IP, II (genes in the interior in both resting and activated Jurkats) and PP (genes at the periphery in both conditions) regions. Statistical significance was determined by Fisher tests. \**P* < 0.05, \*\**P* < 0.01 and \*\*\**P* < 0.001. (*D*) Genome browser views for the genomic region surrounding the GBP gene cluster and *CBLB, CD200, BTLA* and *IL2* loci covering the same parameters as in B. Each gene is shaded in light green over the various traces. (*E*,*F*) Representative micrographs and quantification of gene positions in resting and activated cells relative to lamin B2. The median points are marked with dashed red lines. For quantification statistics, the position of loci in the activated sample was compared to the resting sample by KS tests. \**P* < 0.05, \*\**P* < 0.01 and \*\*\**P* < 0.001. See also Supplemental Fig. S4.

Accordingly, PI regions released from the periphery contained significantly more induced than repressed genes (Fig. 3C). In the opposite direction, IP regions moving to the periphery displayed significantly more repressed than induced genes (Fig. 3C). Similarly, like IP regions, loci remaining at the periphery (PP genes) displayed a significant enrichment of repressed genes. Nonetheless, despite the association of the periphery with transcriptional repression, the majority of genes moving to or from the nuclear periphery displayed no change in expression (Supplemental Fig. S3D). Interestingly, while the trend towards the periphery being a repressive environment was also observed in a recent study of myogenesis (Supplemental Fig. S3C) (Robson et al. 2016), the association between repositioning and altered gene expression was much less in the lymphocyte activation model studied here (Supplemental Fig. S3, compare panels D and E). Processed DamID and microarray data for all genes can be found in Supplemental Table S1.

The subset of genes undergoing both repositioning and expression changes tended to be important for T-cell regulation based on their overlap with the aforementioned lymphocyte activation functional classes and GO-terms associated with lymphocyte function. To confirm their decreased peripheral association inferred from DamID, five genes were selected for direct testing by fluorescence *in situ* hybridization (FISH). These genes were chosen based on their being induced throughout the Jurkat T-cell activation time course (Fig. 3D, Expr. tracks) and also being present in PI regions (Fig. 3D, log2(B1/Dam) IP/PI region tracks). These were the guanylate binding protein (*GBP*) gene cluster, Cbl proto-oncogene B (*CBLB*), CD200 molecule (*CD200*), B and T lymphocyte associated (*BTLA*) and interleukin 2 (*IL2*). GBP1 regulates the cytoskeleton during T-cell activation (Sharma et al. 2011) while IL-2 is a potent inducer of T-cells (Boyman and Sprent 2012). The other three genes chosen appear more focused on regulating or attenuating immune responses with *CBLB* encoding a ubiquitin ligase and *BTLA* and *CD200* respectively encoding a inhibitory ligand and receptor (Watanabe et al. 2003; Rygiel et al. 2009; Wallner et al. 2012). FISH was performed in resting and activated Jurkat T-cells together with lamin B2 staining to identify the edge of the nucleus. The absolute distance between the peak fluorescence intensity of lamin B2 and the gene was measured in the mid-plane from deconvolved wide-field images and plotted as a cumulative frequency plot. In such plots increasing distances from the lamina are viewed as a shift to the right. The directly measured gene positions for the *GBP* cluster, *CBLB, CD200, IL2* and *BTLA* all matched the peripheral positioning predicted by the DamID in resting cells and were released during activation (Fig. 3E, F). In all cases the differences were highly statistically significant by Kolmogorov-Smirnov test. By contrast, an activated PP gene (*i.e*. present in LADs in both resting and activated cells) that is important for T-cell associations with the vascular endothelium termed vascular cell adhesion molecule 1 (*VCAM1*), did not display an increased distance to the periphery following T-cell activation (Supplemental Fig. S4A-B). This supports our methodology and confirms that not all genes reposition away from the periphery during activation. As an additional control, peripheral distance measurements were also performed for *IL2RA/CD25*, the gene encoding the high-affinity IL-2 receptor subunit alpha that is induced in T-cell activation and lacks lamina-association in resting or activated cells (Supplemental Fig. S4C, D). Consistent with the DamID data, the *IL2RA* locus is significantly further from the nuclear envelope than lamina-associated genes. Moreover, following T-cell activation no increase in distance between the *IL2RA* locus and the nuclear envelope was observed. The absence of a change in distance from the periphery measured for *IL2RA* also serves as a control that the increases measured for the *GBP* cluster, *CBLB, CD200, BTLA* and *IL2* loci represent bona fide gene repositioning rather than reflecting global increases in nuclear volume.

### Peripherally sequestered enhancers are released from the periphery upon their induction

Enhancers, *cis*-regulatory elements that activate target genes by physical proximity, become transcribed prior to their functional activation (Arner et al. 2015). As the degree of gene activity correlated with the frequency of lamina-association, we predicted that enhancers, like genes, would reposition away from the periphery concomitantly with their activation during T-cell activation. To test this, a list of enhancer clusters/super-enhancers identified as functioning during T-cell activation by their accumulation of H3K27ac were extracted from a previous study (Hnisz et al. 2013) and compared to the DamID data for Jurkat activation. Of the nearly 1,000 enhancer clusters, 354 were active in both states, 107 were active only in resting T-cells, and 513 were induced only in activated T-cells (Fig. 4A). Of those induced during T-cell activation, only 27 occurred in LADs in the resting Jurkat T-cells. However, over half of these were released from the periphery in PI regions in the activated Jurkat T-cells (Fig. 4B).

**Figure 4.**
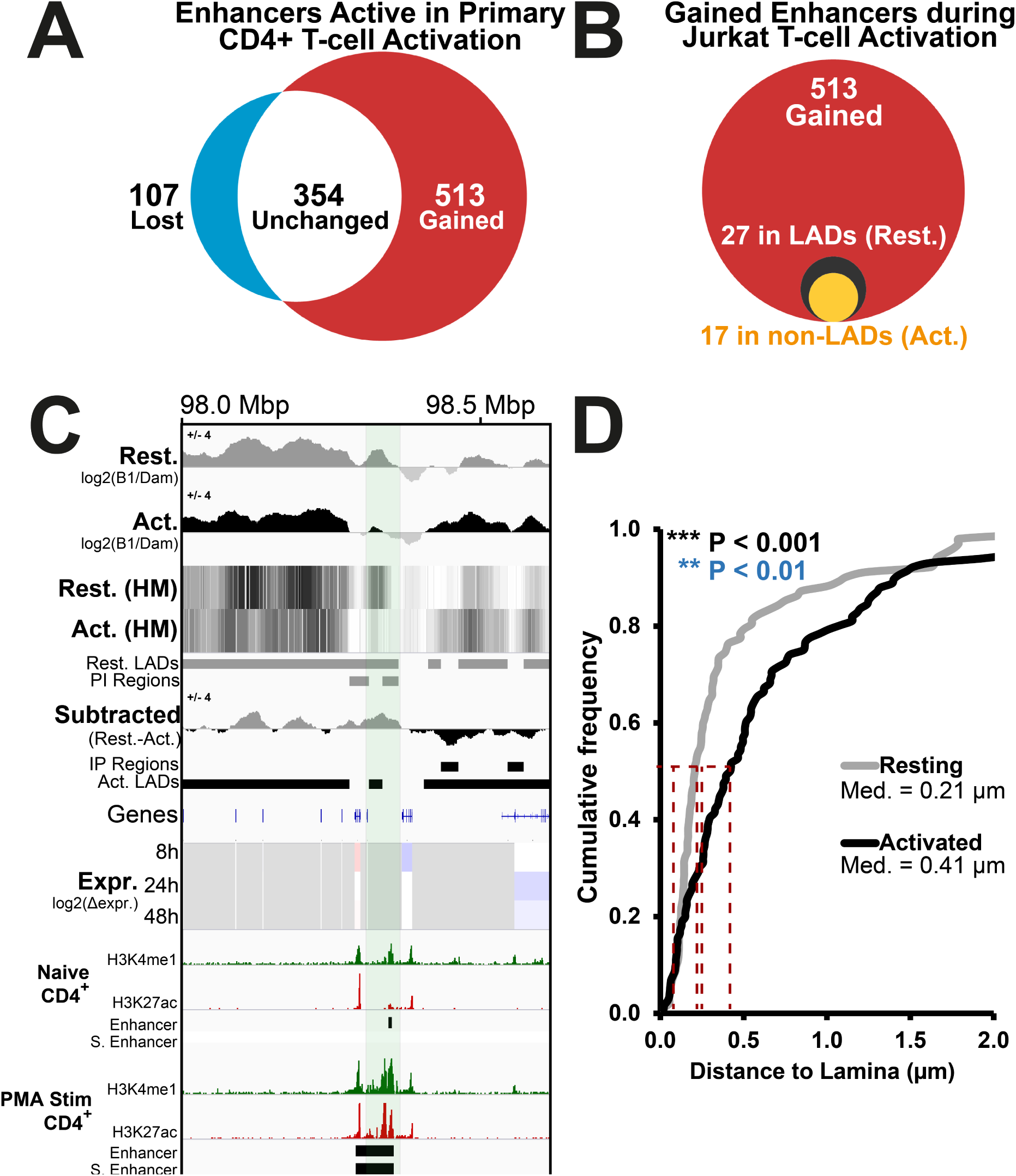
Enhancers are released from the periphery during T-cell activation. (*A*) Upon stimulation of naïve CD4^+^ T-cells with PMA, 513 out of a total of 974 predicted enhancer clusters identified by Hnisz et al, 2013 are induced (red) while only 107 are silenced (blue). (*B*) Of the 513 enhancer clusters that are induced upon activation, most are located in non-LAD regions in the resting Jurkat T-cells, but 27 were in LADs (black circle). Of the 27 enhancer clusters that were associated with LADs in the resting state, 17 become released upon T-cell activation (yellow circle). (*C*) Genome browser view displaying the DamID signal, LADs, IP and PI regions and gene expression changes in Jurkat T-cells for a predicted enhancer on chromosome 3. Published ChIP-seq tracks from primary naïve and PMA-activated CD4^+^ T-cells for H3K4me1 and H3K27ac histone modifications marking active enhancers are also shown (Data taken from Bernstein et al, 2010). A 66 kbp region containing a putative enhancer induced during T-cell activation displays release from the periphery (PI) in activated Jurkat cells. The H3K27ac peaks defined as Enhancer and Super Enhancer (S. Enhancer) by Hnisz et al are also displayed as tracks. (*D*) Quantification of putative enhancer position in resting and activated Jurkat T-cells cells relative to lamin B2, confirming its release from the periphery following Jurkat T-cell activation as in Fig. 3. For quantification statistics, the position of loci in the activated sample was compared to the resting sample by KS tests. \**P* < 0.05, \*\**P* < 0.01 and \*\*\**P* < 0.001.

To confirm the peripheral release of activated enhancers, FISH was performed on an enhancer cluster devoid of genes which displayed decreased peripheral association in stimulated Jurkat T-cells and accumulated H3K27ac and H3K4me1 upon activation of human CD4^+^ T-cells with Phorbol 12-myristate 13-acetate (PMA) (Bernstein et al. 2010) (Fig. 4C). As predicted, the distance between the enhancer locus and the periphery increased following its induction during T-cell activation (Fig. 4D). Hence, similar to genes, peripherally positioned enhancers are released concomitantly with their activation and acquisition of H3K27ac during T-cell activation.

### Topologically Associated Domains form the unit of gene repositioning

LADs have been reported to significantly overlap with TADs, regions of chromosomes that display preferential local interactions (Dixon et al. 2012; Fraser et al. 2015; Kind et al. 2015; Jabbari and Bernardi 2017). As TADs have been shown to be largely invariant between different cell types (Rao et al. 2014; Dixon et al. 2015; Fraser et al. 2015), we hypothesized that regions with altered peripheral localization would be delimited to single TADs rather than spanning multiple TADs.

To investigate this we contrasted the DamID maps of resting and activated Jurkat T-cells to the published GM12878 lymphoblastoid cell line Hi-C dataset (Rao et al. 2014). Notably, in this study regions of preferential local interactions were defined using a different methodology than historically used to define TADs, though they were largely similar (Rao et al. 2014). As we use the exact coordinates for the “Contact Domains” that they determined in their published study, we use the Rao et al terminology rather than referring to these structures as TADs.

Global analysis of Contact Domains and LAD distributions between the GM12878 Hi-C map and our DamID dataset for resting and activated Jurkat T-cells revealed that LADs are frequently contained in their entirety within Contact Domains. This was the case for both resting and activated LADs (Supplemental Fig. S5A). To test this possibility 500 LADs were randomly selected and tested for their complete envelopment within Contact Domains. As a control, frequency of complete envelopment within Contact Domains was then also measured for the same 500 LADs after they had been randomly shuffled along their chromosome. Significantly, non-mappable regions such as centromeres were excluded from the LAD shuffling. This process was then repeated over 1,000 iterations to ensure all LADs in the genome were analyzed multiple times over the total sampling and the frequencies of LAD envelopment within Contact Domains plotted (Supplemental Fig. S5). Both resting and activated LADs occurred in their entirety within Contact Domains significantly more frequently than for the shuffled genome iterations while no statistically significant difference was observed between the resting and activated LADs. We postulate that this observed increase in the expected frequencies of LADs found completely within Contact Domains indicates that when a Contact Domain contains a region of altered peripheral association the Contact Domain repositions as a unit. A similar analysis was also performed using the GenometriCorr package, a tool that performs multiple analyses testing the spatial independence of domains (Supplemental Fig. S5B) (Favorov et al. 2012). Confirming the previous analysis, the GenometriCorr testing indicated the positions of Contact Domains and TADs were highly correlated, displayed a uniform distance separation and overlapped significantly more than would be expected by random chance.

To test that Contact Domain structure is similar in GM12878 cells and Jurkat T-cells, we chose 5 fosmid probes containing sequences that were separated by 149 kbp from each other in series over the *VCAM1* locus (Fig. 5A). The first was in a neighboring Contact Domain upstream of *VCAM1 (i)*, the second just within the *VCAM1* Contact Domain boundary *(ii)*, the third directly in the Contact Domain center *(iii)*, the fourth just within the Contact Domain boundary downstream of *VCAM1 (iv)*, and the fifth downstream in a neighboring TAD *(v)*. The distance between three different equally spaced probe pairs, two crossing a TAD boundary (pair 1:i+iii and pair 3:iii+v) and one contained within the same Contact Domain (pair 2:ii+iv), was determined in 2D from max projection deconvolved images taken by widefield fluorescence microscopy. Supporting the similarity between the Contact Domain structure in GM12878 cells and resting Jurkat T-cells, the probes within the same Contact Domain were significantly closer (0.15 μm) than the probes between adjacent Contact Domains (∼0.30 μm) (Fig. 5B). The depth and quality of the GM12878 data is underscored by its validation by direct testing, whereas if a TAD definition was used (albeit from human embryonic stem cells Hi-C data; Dixon et al, 2012) probe pairs 1 and 2 should have had a similar separation distance.

**Figure 5.**
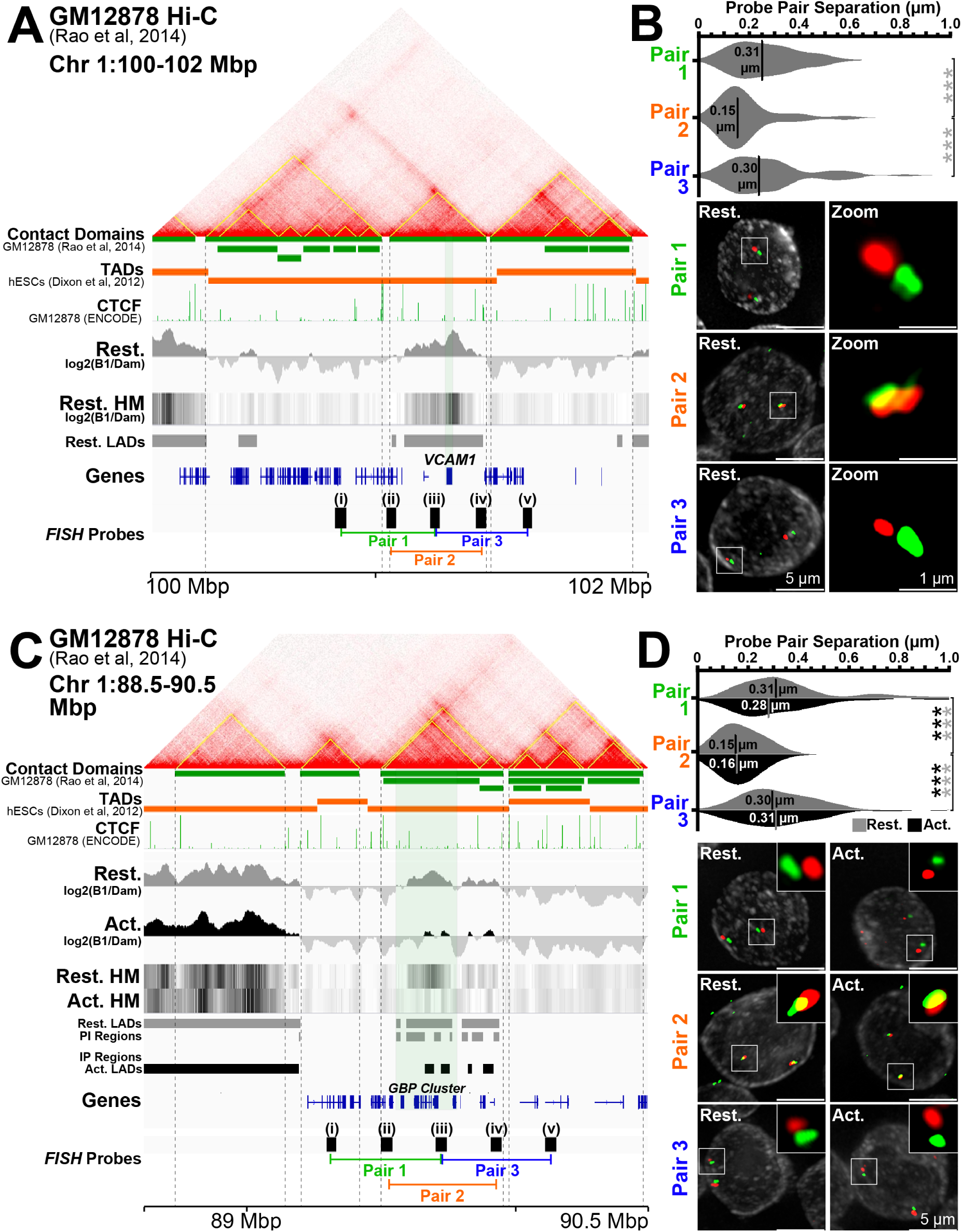
The unit of repositioning is the Contact Domain. (*A*) Genome browser view for the genomic region surrounding *VCAM1* showing Hi-C data and associated Contact Domains (yellow lines on contact frequency heatmap and green maps in genome browswer track) from published data determined in GM12878 lymphoblastoid cells (Rao et al. 2014). Directly under the contact domains are shown TADs (orange lines) from published data determined in human embryonic stem cells (hESCs) (Dixon et al. 2012). Being important for the establishment of Contact Domains/TADs, a published ChIP-seq track for CTCF in GM12878 cells is also shown (ENCODE Consortium, 2012). DamID signal intensities, identified LADs, IP and PI regions and microarray gene expression changes for resting and activated Jurkat T-cells are shown below. In addition shown below are probes and probe pairings tested in by FISH in panel B. (*B*) Quantification and representative max projection images displaying distance between indicated sets of ∼150 kbp spaced FISH probe pairs. Pairs 1 and 3, containing probe pairings from different Contact Domains, are separated significantly further than probes within the same Contact Domain used for Pair 2. (*C*) Identical genome browser views of the *GBP* gene cluster and probes tested. (*D*) Quantification and representative images displaying distance between indicated sets of ∼175 kbp spaced FISH probe pairs in resting and activated Jurkat T-cells. While Pairs 1 and 3 are again separated more than Pair 2, the distances separating the probes in all pairs remain unchanged during T-cell activation indicating the Contact Domain structure is maintained. Quantification statistics were determined by comparing the separation of Pairs 1 and 3 with Pair 1 in resting (grey) and activated (black) cells by KS tests. \**P* < 0.05, \*\**P* < 0.01 and \*\*\**P* < 0.001. See also Supplemental Fig. S5.

To test whether a Contact Domain is the unit of repositioning or if its structure changes during lymphocyte activation-induced gene repositioning, FISH was performed on resting and activated Jurkat T-cells using a similarly generated set of 5 FISH probes (except 174 kbp distant from one another) covering the GBP gene cluster (Fig. 5C). Again, a significantly reduced distance was observed between probe pairs within the Contact Domain than between adjacent Contact Domains and this organization was maintained between both the resting and the activated states (Fig. 5D). Hence, at least in this instance, the unit of repositioning is the Contact Domain and it remains unaltered, at least at this resolution, following release from the lamina.

### Specific chromosome compartments segregate with respect to LADs

Compartments represent interactions between regions of chromosomes that share a similar functional state, contrasting the proximal local interactions defining TADs/Contact Domains (Lieberman-Aiden et al. 2009; Dixon et al. 2012; Nora et al. 2012; Rao et al. 2014). These functional states are broken into transcriptionally active A compartments and inactive B compartments that, in the high-resolution GM12878 cell data, can each be further subdivided (Rao et al. 2014). Strikingly, the B sub-compartments, defined solely from Hi-C interactions, each display distinct heterochromatin states, B1 being significantly enriched in Polycomb H3K27me3 and B2 and B3 correlating with LADs (Lieberman-Aiden et al. 2009; Gibcus and Dekker 2013; Rao et al. 2014). We do not consider B4 sub-compartments as they only represent a small fraction of the genome restricted to chromosome 19. By contrast, the A sub-compartments are very similar to one another in the chromatin states measured, with A2 displaying increased H3K9me3 (Rao et al.2014). As we now have mapped regions specifically changing at the periphery between resting and activated Jurkat cells, we sought to determine if the Jurkat LADs and IP/PI regions correlated with any particular sub-compartment from the GM12878 data.

First, we extracted the Rao et al sub-compartment regions and separately 1kb upstream and downstream flanking DNA. As they vary in size, all individual compartment regions were then normalized to an equal length in the plot. The average Log2(Lamin B1/Dam-only) ratio from resting Jurkat T-cells was then plotted over these size-normalized regions and the 1 kb regions of upstream and downstream flanking DNA (Fig. 6A). As anticipated, in the resting Jurkat cells the B2 and B3 sub-compartments displayed a peak of Log2(Lamin B1/Dam-only) signal relative to flanking regions, while the A1 and A2 sub-compartments displayed a depleted Log2(Lamin B1/Dam-only) signal. By contrast, the polycomb-associated B1 sub-compartment displayed only a modest increase in Log2(Lamin B1/Dam-only) signal relative to the flanking DNA. Accordingly, we observe LADs as significantly enriched in B2 and B3 compartments while being significantly depleted in B1 and A1 sub-compartments relative to a genome where LADs had been randomly shuffled (Fig. 6B). This further corresponded with their expression, A1 sub-compartment being the most highly expressed followed by A2, B1 and B2/B3 sub-compartments (Fig. 6C).

**Figure 6.**
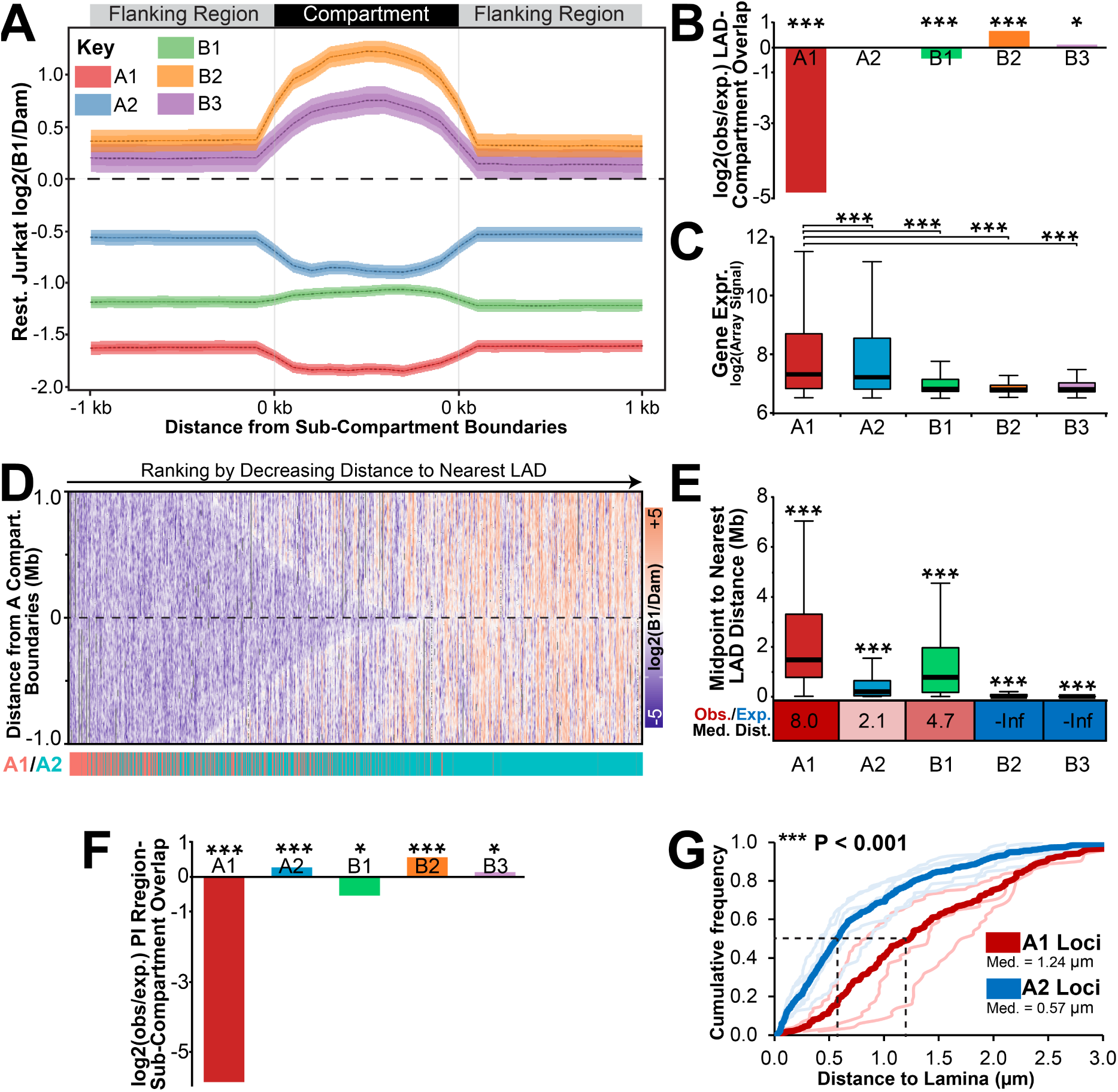
Compartments are distributed non-randomly with respect to LADs. (*A*) Size normalized plot displaying the average log2(Lamin B1/Dam-only) signal from resting Jurkat T-cells across all A1, A2, B1, B2 or B3 GM12878 sub-compartment regions and 1 kbp of flanking DNA. Compartment co-ordinates were extracted from Rao et al, 2014. In order to compare all regions of a given sub-compartment, region sizes were normalized to a uniform standard. (*B*) Fold enrichment analysis of LADs observed (obs) in each compartment compared to that expected in a genome with randomly shuffled LADs (exp). (*C*) Box plot displaying the distribution of levels of gene expression in resting Jurkat T-cells for loci in GM12878 sub-compartments. (*D*) Heatmap of DamID log2(B1/Dam) signal for +/-1 Mbp surrounding all A compartment regions. Regions have been ranked by decreasing distance to nearest LADs and their identity as an A1 or A2 subcompartment region highlighted below in red or blue, respectively. A2 domains are more observed more frequently in regions more proximal to LADs. (*E*) Box plot displaying the distribution of chromosomal distances between the midpoint of sub-compartment regions and the nearest LAD in resting Jurkat T-cells. Below is the ratio between observed (Obs.) median distance and that expected (Exp.) in a randomly LAD-shuffled genome. Values have been colour coded to show increased (red) or decreased (blue) observed distance compared to expected. (*F*) Fold enrichment analysis of PI regions observed in each compartment compared to that expected in a genome with randomly shuffled PI regions. (*G*). Quantification of average distance between A1 (thick orange line) or A2 loci (thick grey line) and the lamina determined by FISH. A2 loci included *GBP, NFKBIZ, CBLB, CD200, BTLA* (as displayed in Fig. 3E) while A1 loci included *IL2RA* (as displayed in Fig. S4A), *CCND2* and *TNF* and are displayed individually as lighter, thinner lines below. FISH analysis was performed in activated Jurkat T-cells cells, *i*.*e*. where all loci measured are known to be detached from the nuclear lamina. For B and F significance was determined by Fisher tests while for C and E significance was by Dunn test for multiple comparison testing after a Kruskal Wallis significance test. For G the significance of the difference between A1 and A2 distances to lamina was determined by KS tests. \**P* < 0.05, \*\**P* < 0.01 and \*\*\**P* < 0.001. See also Supplemental Fig. S6.

The DamID signal was higher in the regions flanking B2 and B3 sub-compartments than the regions flanking the other compartments (Fig. 6A). Hence, the B2 and B3 sub-compartments are not only more likely to be in a LAD but also to be flanked by LADs. Interestingly, the next highest log2(Lamin B1/Dam-only) intensities were associated with the A2 sub-compartment and their flanking DNA, roughly three times higher than the A1 sub-compartment. As there was little difference in A1 and A2 compartments in previous analyses (Rao et al. 2014), this striking difference in flanking lamina-association suggested to us that these represent spatially segregated compartments with respect to the nuclear periphery.

We plotted the Log2(Lamin B1/Dam-only) signal within 1 Mbp of all A sub-compartment region boundaries and ranked them based on their distance to the nearest LAD. Surprisingly, this heatmap revealed that the A1 sub-compartment regions tended to be further from LADs than the A2 sub-compartment regions (Fig. 6D). To quantitate this distinct distribution of A1 and A2 regions, we then plotted the distances from the midpoint of sub-compartment regions to the nearest LAD in the observed data (Fig. 6E). This revealed, as expected, that B2 and B3 sub-compartment regions were very close to LADs (with a median distance of 0 bp to the nearest LAD), much closer than would be expected in a genome where LADs had been randomly shuffled. The A2 subcompartment regions were next closest to LADs (median distance 193 kbp) while B1 and A1 subcompartments were significantly further away (median distances respectively of 767 kbp and 1.468 Mbp). Supporting the fact that sub-compartments are less frequently in LADs, A1, A2 and B1 subcompartments all were further away from LADs than expected in a genome with randomly shuffled LADs (Fig. 6E). Similar statistically significant overlaps, or lack thereof, of sub-compartments with LADs was also observed by GenometriCorr analysis (Supplemental Fig. S6A). Therefore, we propose, due to difference in their proximity to LADs along chromosomes, that the A1 and A2 subcompartments represent active chromatin that is spatially segregated with respect to the periphery.

It follows that a gene being released from the periphery during lymphocyte activation would be more likely to enter an A2 than an A1 sub-compartment. Unfortunately, the Hi-C data we used from the Rao study was not matched to our resting and activated DamID datasets and moreover the activation state of their cells was not clearly established. However, a comparison of the Jurkat T-cell resting and activated DamID data with the Rao et al. 2014 GM12878 compartment maps revealed that several of the genes we had directly tested that were at the periphery in resting Jurkat T-cells and released in the activated appeared in the A2 compartment in the GM12878 data. Similarly, just as in activated T-cells, PI genes displayed elevated expression in GM12878 cells relative to resting Jurkat T-cells (Supplemental Fig. S6B). Therefore we proceeded on the assumption that the GM12878 cells were in a state more reflective of activated than resting Jurkat T-cells. Thus, different from the GM12878 Rao et al, 2014 data, the presence of these genes in LADs in the resting Jurkat T-cells would reflect a different state in which they are in a B2/B3 subcompartment until T-cell activation where they are induced and released to associate preferentially with the A2 sub-compartment. Indeed, we observe that PI regions are significantly enriched in the A2 sub-compartment in GM12878 (Fig. 6F) cells and have a high content of activated genes relative to resting Jurkats (Fig S6B). Similar results were observed using GenometriCorr analysis, except that the B1 compartment was also observed to be enriched in PI regions (Fig S6C). Moreover, supporting the A2 compartments restriction near the lamina, genes that were both in A2 compartments in GM12878 cells and released from the periphery and induced in activated T-cells were significantly closer to the lamina than three A1 sub-compartment loci tested (Fig. 6G).

### Release of a predicted T-cell activation-specific enhancer from the nuclear periphery enables its association with similarly released genes in the interior

The Jurkat activation system further enabled direct testing of this repositioning to a proximal compartment by FISH as it would be predicted loci which are released from a LAD B compartment to a lamin proximal A2 compartment during T-cell activation would display increased proximity in stimulated T-cells. Therefore, a 16 Mbp region of chromosome 3 was selected where several Jurkat activation PI genes and a predicted enhancer corresponded to a compartment identified in the Rao study (Fig. 7A). Several of these genes were tested in Figure 3 to confirm release from the periphery concomitant with their induction during Jurkat T-cell activation (*CBLB, BTLA* and *CD200*). We also tested the positioning of the *NFKBIZ* locus, which was induced during T-cell activation, also found in an A2 sub-compartment in GM12878 cells and was proximal to but not immediately overlapping with a PI region. Though only immediately proximal to a LAD, this locus was close to the periphery in resting cells and became more internal during activation, indicating again that genomic context is important to consider in addition to the LADs themselves (Fig. 7B).

**Figure 7.**
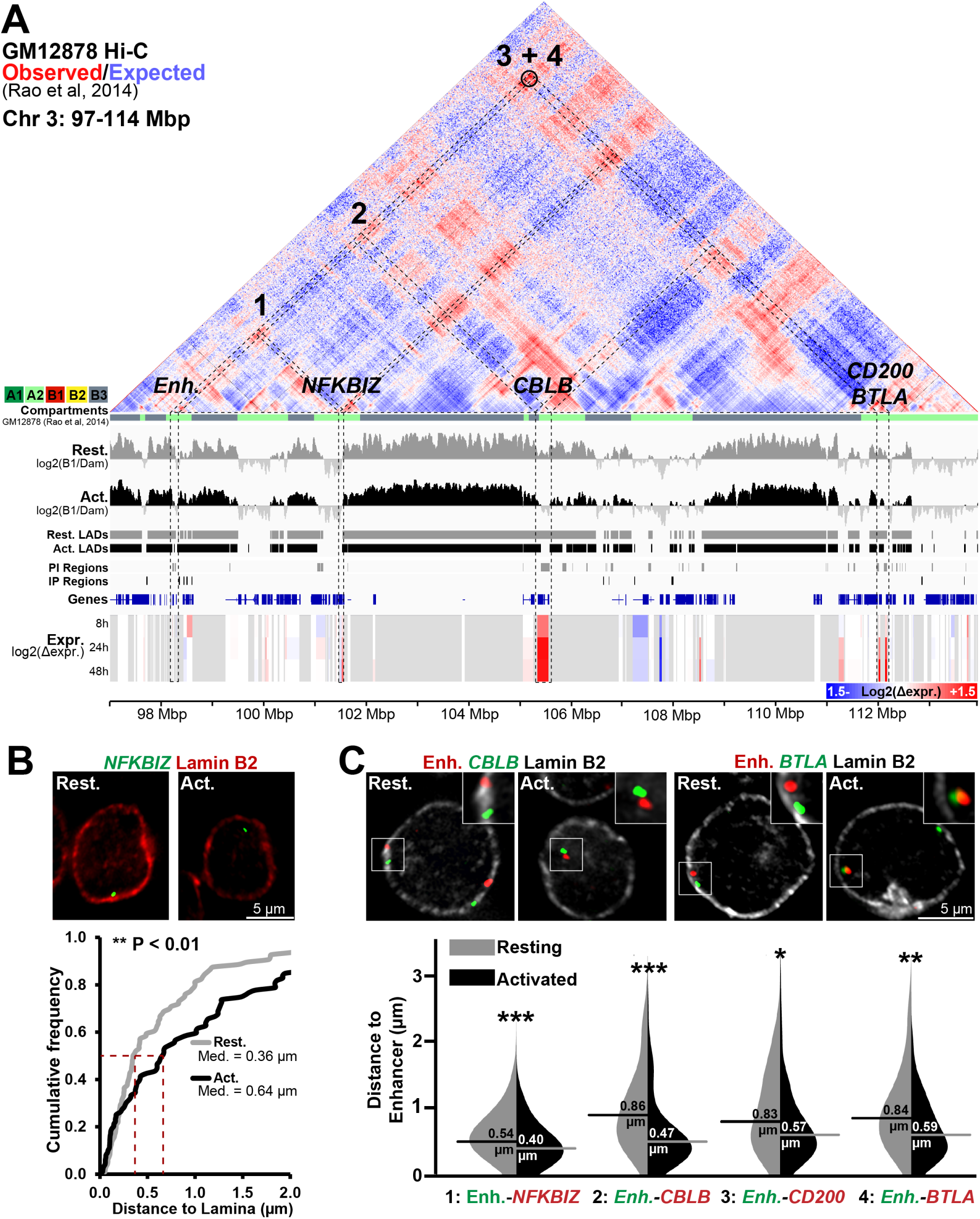
Genes released from LADs during T-cell activation associate in the interior in activated Jurkat cells. (*A*) Genome browser view of a 17 Mbp region of chromosome 3 displaying GM12878 cell Hi-C interactions from published contact maps (Rao et al. 2014). Hi-C data is displayed as a heat map of the ratio of observed contact frequencies over that expected in a distance scaled model. GM12878 sub-compartments (A1, A2, B1, B2, B3) are color coded directly beneath the contact data. DamID signal intensities, identified LADs, IP and PI regions and microarray gene expression changes for Resting and Activated Jurkat T-cells are also displayed as in previous figures. The blue circle represents highly significant DNA-DNA contacted detected by Rao, et al. Several genes, *NFKBIZ, CBLB, CD200* and *BTLA*, and a T-cell activation-specific enhancer (Enh.), display reduced lamina association during T-cell activation concomitant with transcriptional induction and are present in the A2 compartment of the GM12878 cells. (*B*) Representative micrographs and quantification *NFKBIZ* positions in resting and activated cells relative to lamin B2. (*C*) Representative micrographs and Violin plot quantification of distances between the enhancer and indicated genes in resting and activated Jurkat T-cells. All enhancer-gene FISH probe combinations display a reduced inter-probe distance following activation. See also Figure S3. For quantification statistics, relative distance measurements between loci or between loci and lamin B2 was compared between the in the activated and resting samples by KS tests. \**P* < 0.05, \*\**P* < 0.01 and \*\*\**P* < 0.001. See also Supplemental Fig. S7.

All these genes were compared to a single locus containing the predicted enhancer described in Figure 4C-D that is released from the periphery upon Jurkat T-cell activation and occurs in an A2 sub-compartment in GM12878 cells. FISH was performed on combinations of the putative enhancer probe with the individual gene probes (*NFKBIZ, CBLB, CD200*, and *BTLA*) in both resting and activated cells. The distance between the loci pairs was then measured using the center of their maximum intensities in 2D images generated from max projections of 3D deconvolved z-series images. For all combinations the distance between the putative enhancer and *NFKBIZ, CBLB, CD200*, and *BTLA* was much greater when the loci were at the periphery in the resting cells than after their release upon activation (Fig. 7C). This is consistent with the idea that they are released from B2/B3 sub-compartments in resting Jurkat T-cells to associate in an A2 sub-compartment. By contrast, the *GBP* cluster and *VCAM1* loci are similarly spaced along chromosome 1 (11 Mbp apart), both are in LADs in resting cells though only *GBP* gets released upon Jurkat T-cell activation, yet no significant change in 3D spatial distance was observed upon activation (Supplemental Fig. S7). Unsurprisingly, the *GBP* cluster and *VCAM1* locus display no Hi-C interactions from the GM12878 data. Likewise, the *IL2* and *IL2RA/CD25* loci that are located on separate chromosomes did not exhibit any significant change in the distance between them (Supplemental Fig. S7B-C). Thus, only when all loci activated within a chromosomal region were released from LADs did they display increased spatial proximity in the nuclear interior following T-cell activation.

### The release and association of the predicted enhancer with *CBLB, CD200*, and *BTLA* also occurs in primary lymphocyte activation

The DamID was necessarily performed in the Jurkat model T-cell activation system due to poor transduction efficiencies of primary lymphocytes and the possibility of activation during lentiviral treatment. However, the differences in the activation state of a small subset of the lymphocyte activation gene classes (Fig. 2 and Supplemental Fig. S2) necessitated validation of these findings in primary human lymphocytes.

A population of naïve CD4^+^ T-cells was isolated from fresh human peripheral blood mononuclear cells (PBMCs) by polychromatic FACS (Supplemental Fig. S8). Part of this population was directly fixed for FISH while another was activated with PMA for 24 h. PMA was chosen for activation to better match results to the published ChIP-Seq study where lymphocytes were also activated with PMA (Bernstein et al. 2010). A portion of the activated cells was fixed for FISH while the level of activation was measured on another portion by the presence of CD69+ by FACS analysis (78%; Fig. 8A). The fixed cells were hybridized with the same probe pairs used in Figure 7. In all cases the predicted enhancer became closer to the gene locus and this was significant for all except for *NFKBIZ* (Fig. 8B). Also, as expected, the predicted enhancer, *NFKBIZ, CBLB, CD200*, and *BTLA* were all at the periphery in the resting primary lymphocytes and all moved away upon activation (Fig. 8C). In all cases this repositioning was highly significant. Thus, these findings accurately reflect genome organization changes that occur upon human lymphocyte activation at least for the genes tested.

**Figure 8.**
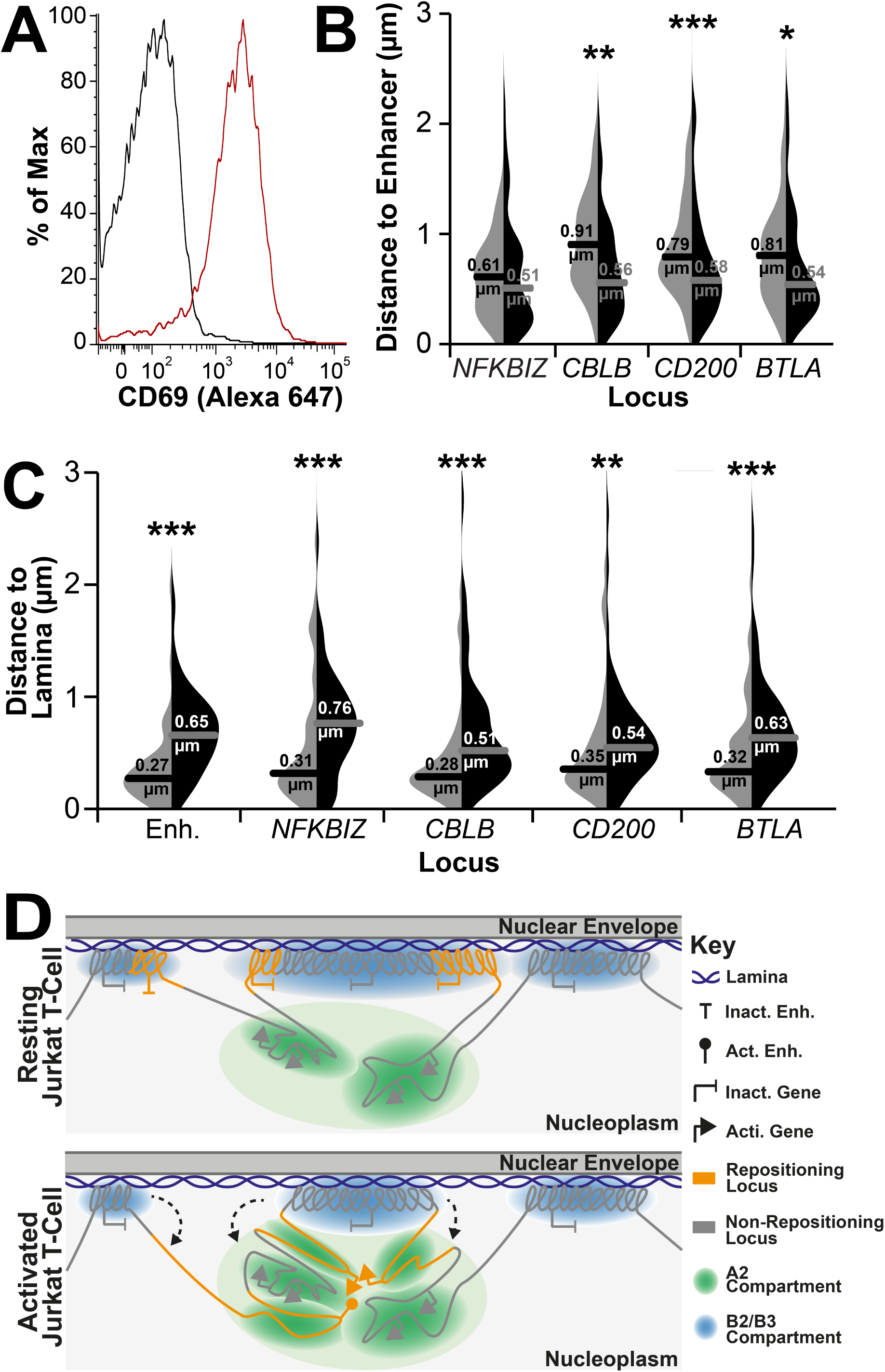
LAD release and association of genes in the nuclear interior occurs in primary CD^4+^ T-cells. (*A*). FACS plot of CD69 staining for naïve (resting, black line) and PMA-treated (activated, red line) CD4^+^ T-cells indicating ∼78% cell activation. See also Supplemental Fig. S8 for details of the isolation of the naïve CD4^+^ T-cell population. (*B*) Violin plot quantification of distances between the predicted enhancer and indicated genes in naïve (light grey side) and PMA-treated (black side) cells. All loci displayed closer proximity to the predicted enhancer locus upon PMA activation of naïve CD4^+^ T-cells. (*C*) Violin plot quantification of distances between indicated loci and lamin B2 in naïve (light grey side) and PMA-treated (black side) cells. All loci displayed increased distance from the periphery upon PMA activation of naïve CD4^+^ T-cells. For quantification statistics, relative distance measurements between loci or between loci and lamin B2 was compared between the in the activated and resting samples by KS tests. \**P* < 0.05, \*\**P* < 0.01 and \*\*\**P* < 0.001. (*D*) Schematic model of the relationship between LADs, compartments and gene release. In resting T-cells B2 and B3 sub-compartments exist as LADs while A2 sub-compartment regions remain near to the lamina due to their proximity along the chromosome. By contrast the A2 and B1 compartments, being further from LADs along chromosomes, are distributed more distally in the nuclear interior (not shown). Following T-cell activation, regions of LADs containing necessary gene or enhancer loci are released from the periphery where they associate in the A2 compartment proximal to the lamina in a transcriptionally active state.

## Discussion

### Genes subject to spatial regulation are important for T-cell activation

In an earlier study of myogenesis the genes specifically regulated from the nuclear envelope were often required in the initial stages of differentiation, but needed to be tightly shut off later because they were inhibitory at later stages (Robson et al. 2016). Similarly, here, while first responder genes that initiate activation were strongly absent from LADs, the genes subject to spatial regulation were important for subsequent T-cell function and regulation. For example, *IL2* encodes the cytokine interleukin 2, a potent inducer of subsequent T-cell activity and function in propagating an immune response (Boyman and Sprent 2012), while *GBP1* encodes the GTPase guanylate binding protein 1 which regulates the cytoskeleton to influence T-cell activation signalling (Sharma et al. 2011). If these activities were not tightly shut down prior to immune activation they could potentially activate an immune response in the absence of infection.

Likewise, several of the spatially regulated genes associated with this putative enhancer function in modulating the activation response. For example, induction of the nuclear IκB family member encoded by *NFKBIZ* during T-cell activation is required for the expression of *IL17* following T-cell stimulation (Okamoto et al. 2010). Similarly, *BTLA* and CD200, encoding an inhibitory receptor and ligand respectively, and *CBLB*, encoding a E3 ubiquitin ligase, are expressed following T-cell stimulation and act to attenuate T-cell activation and thus fine tune the immune response (Watanabe et al. 2003; Rygiel et al. 2009; Wallner et al. 2012). As a result of these important activities in T-cell regulation, the dysregulation or loss of function of a number these peripherally regulated gene products is associated with autoimmunity (Watanabe et al. 2003; Rosenblum et al. 2004; Sharma et al. 2011; Wallner et al. 2012; Johansen et al. 2015; Hussman et al. 2016). Thus in all cases the tighter regulation of these genes in resting lymphocytes from adding spatial control would benefit the organism. Similarly regulating *cis*-regulatory elements such as enhancers is likely beneficial due to their ability to act on multiple target genes (Symmons and Spitz 2013).

### The relationship between gene activity and lamina association

It is clear from this study that specific gene repositioning both to and from the periphery occurs during lymphocyte activation, but it remains unclear what controls the initial tethering or subsequent release of loci. Our analysis indicates that the frequency of lamina-association was inversely related to the level of gene expression across all global metrics investigated. This is consistent both with previous work globally measuring peripheral position and expression (Guelen et al. 2008; Peric-Hupkes et al. 2010; Akhtar et al. 2013; Robson et al. 2016) and with artificial targeting of transcriptional activation to release a locus from the periphery (Tumbar et al. 1999; Tumbar and Belmont 2001; Chuang et al. 2006; Therizols et al. 2014). Thus, it is possible that transcriptional activation drives the release of genes from the periphery upon stimulation of T-cell activation. The same could apply for enhancers that are frequently transcribed during their induction (Arner et al. 2015). However, it is important to note that the relationship between gene position and expression is not absolute as the activity of the majority of repositioning genes remains unaffected during T-cell activation (here), during myogenesis (Robson et al. 2016) and during neurogenesis (Peric-Hupkes et al. 2010).

Chromatin unfolding induced by artificial targeting of an acidic peptide also caused gene release in the absence of transcription, indicating chromatin state rather gene activity per se, regulates lamina association (Therizols et al. 2014). Supporting this, depletion of repressive chromatin modifications enriched in LAD bodies or borders, such as H3K9me2/3 or H3K27me3, respectively, results in disrupted lamina association (Zullo et al. 2012; Bian et al. 2013; Demmerle et al. 2013; Kind et al. 2013; Harr et al. 2015). These studies suggest that the marks not only provide for silencing, but themselves contribute to locus affinity for the periphery. At the same time, loci can be recruited to the nuclear periphery with an artificial tether (Finlan et al. 2008; Reddy et al. 2008) and specific nuclear envelope proteins have been shown to be important for peripheral tethering of endogenous developmentally repositioning loci (Zullo et al. 2012; Solovei et al. 2013; Robson et al. 2016). Several studies argue that the physical recruitment of a locus to the periphery alone is insufficient to silence the locus, but works together with transcriptional regulators and other silencing factors at the periphery (Finlan et al. 2008; Robson et al. 2016). Once at the periphery, other nuclear envelope proteins that have been shown to recruit enzymes, such as histone deacetylases, that add epigenetic marks can reinforce a repressed state (Somech et al. 2005; Demmerle et al. 2012). Thus, tethering to the nuclear envelope is a function of the chromatin state together with the milieu of nuclear envelope proteins. Consequently, tethering interactions, repressive marks, chromatin folding state and transcriptional activation likely all compete for dominance until a threshold is achieved for release. In this regard, release of enhancers could enable a step-wise alteration of gene expression due to the fact that gene activity would not reach optimal levels until the enhancer was also released.

### Lamina-association provides a compass for interpreting compartment information

By contrasting both Hi-C data and LADs we now place lamina-association and gene release within the context of higher order chromatin structure. Chromosome compartments are defined based on contact frequencies from Hi-C data and accordingly reveal what is close to each other, but not where the compartment is located within the nucleus. Previous studies comparing Hi-C and DamID datasets revealed a tendency for B compartments to be enriched in LADs, nucleolar-associated domains (NADs) and H3K27me3 Polycomb, both of which represent spatially segregated heterochromatin (Lieberman-Aiden et al. 2009; Gibcus and Dekker 2013; Rao et al. 2014). However, no spatial discrimination of active compartments has been previously described. Here we found that A1 and A2 sub-compartments segregate with their proximity to LADs along chromosomes, with the A2 sub-compartment occurring closer to the lamina. Thus the physical anchoring of specific regions in LADs determines the relative spatial positions of the wider genome. It follows that differences observed in DamID data between different cell types and stages of differentiation could have wider consequences for the regulation of genes in the nuclear interior.

Tellingly, the genes released from the nuclear periphery during T-cell activation exhibited a strong tendency to be in A2 sub-compartments, as well as B2 and B3 sub-compartments, in the GM12878 cells. Our results demonstrate that genes get released from LADs and move into closer proximity in the nuclear interior, relative to a particular locus, following T-cell activation. Moreover, the distance from the periphery achieved by these genes following activation was significantly less than loci from the A1 compartment. Together this strongly suggests these genes are switching from a LAD-associated B sub-compartment to a LAD-proximal A2 sub-compartment.

The restriction in distance from the periphery for released loci could contribute to explaining one of the biggest questions in the spatial genome organization field: how compartments are established and how particular genes can find one another within the large 3-dimensional space of the nucleus. In the analysis here, once released, loci remained relatively close to the periphery, rarely moving further than ∼0.6 μm. This is an important limitation because modelling studies indicate that it is unlikely for loci to find each other in a single cell cycle by diffusion alone if they are 10 Mbp apart (Dekker and Mirny 2016). In contrast, a 0.5-0.8 μm space could be sampled by a locus in 1 h, well within the rapid response timeframe for lymphocyte activation and the distances we measured for loci released from the periphery (0.4-0.59 μm). This “constrained release” could thus increase the likelihood of incorporation of a released locus into an A2 compartment (Fig. 8D) and at the same time keep it close enough to the nuclear periphery to enable its sequestration to the periphery once T-cell activation has abated. Indeed, the activities a number of the released genes have in regulating T-cell function suggest such tight control maybe necessary. A second functional consequence of constrained release from the periphery may also be to promote proximity between enhancers and highly distal target genes while at the same time restricting them away from other inappropriate targets. Intriguingly, the putative enhancer described herein displays a long distance Hi-C looping contact with *BTLA* in GM12878 cells suggestive of such a functional association (Fig. 7A; Rao et al., 2014), however it remains untested if this is functionally significant. Nonetheless, from the data presented here it is clear that organization of specific DNA regions in LADs is intrinsically associated with gene regulation and has significant physical and, possibly, functional consequences for the remaining DNA located in the nuclear interior.

## Methods

### Cell culture and transduction

Jurkat clone E6-1 T-cells and Raji B-cells were cultured in RPMI 1640 supplemented with 10% FBS, 100 units/ml penicillin and 100 μg/ml streptomycin. Jurkat and Raji cells were diluted 1:6 and 1:10, respectively, in fresh media every 2-3 days. Centrifugation at 1,000 rpm for 5 min was performed weekly during passaging to remove cellular debris. VSV-G pseudotyped lentivirus’ encoding DamID, GFP-tagged, or pLKO constructs were generated as described in Supplemental Methods. 10 μg/ml protoamine sulphate was added during transduction to enhance efficiency.

### Formation of immune conjugates with SEE-loaded Raji B-cells

Raji-B cells were incubated for 30 minutes at a concentration of 1 million cells/ml in RPMI 1640 media containing 0.5 μg/ml SEE (Toxin Technologies, Florida, EP404). Subsequently, SEE-loaded Raji cells were pelleted at 1,000 rpm for 5 min, washed twice in PBS by further centrifugation and then resuspended in suspension cell growth medium in 15 ml falcon tube. Conjugated Raji cells were then mixed in a 1:10 ratio with specified Jurkat cell lines to a total cell density of 1 million cells/ml in suspension cell growth medium. Samples were then extracted from mixed cultures at specified times for RNA extraction or fixation. For incubations longer than 1 hour, tube lids were loosened to allow gas exchange between cells within the tube and the incubator.

### Preparation of Peripheral Blood Mononuclear Cells

Human peripheral blood mononuclear cells (PBMCs) were isolated from buffy coats supplied by the Scottish National Blood Transfusion Service (SNBTS) with approval of the SNBTS Committee for Governance of Blood & Tissue Samples for Non-Therapeutic Use and local and national ethics requirements. PBMCs were isolated by density gradient centrifugation using Histopaque-1077 (Sigma). Briefly, blood was diluted 2-fold with PBS and 15 ml was carefully layered over 15 ml Histopaque 1077 in a 50ml tube and centrifuged at 400 × *g* for 30 min at room temperature. The opaque interface containing PBMCs was collected into a fresh tube, diluted in PBS and pelleted. The cells were washed three additional times with PBS before resuspension in RPMI 1640, 10% FBS and incubation at 37°C, 5% CO_2_.

### Flow Cytometry and Sorting

The day after isolation cells were incubated with the following fluorochrome conjugated antibodies: CD4-V450, CD62L-FITC, CD3-PE, CD25-PE-Cy7 (BD Biosciences), CD44-APC-Cy7 and CD69-APC (Affymetrix eBioscience). Unbound antibody was washed away and the cells were sorted using a 4 laser FACS Aria IIu flow cytometer (Beckton Dickinson) running BD FACSDiva v6 Software. An electronic acquisition gate was applied to the Forward/Side scatter plot to exclude debris and aggregates from intact material. Subsequent fluorescence gatings sequentially separated out different populations resulting in a final naïve CD4^+^ T cell population, defined as CD3+ CD4^+^ CD62L+ CD25^-^ CD44^-^ CD69^-^. One population of these cells was fixed and processed for FISH while another population was activated with 20 ng/ml PMA (Phorbol 12-myristate 13-acetate) (Sigma). After 24 h activation one population of activated cells was fixed for FISH while another was analyzed by FACS post fixation with 0.5%PFA for 10 min and staining with CD69-APC antibody. The presence of CD69^+^ was determinant of activation with 78% of cells in the population used for experiments being activated.

### Fluorescence *In Situ* Hybridization (FISH)

For FISH experiments 50 μl of Jurkat or primary CD4^+^ T-cells were pelleted and resuspended in PBS and then added to polylysine-coated coverslips and allowed to settle for 15 min. Cells were then fixed with 4% paraformaldehyde (PFA), 1X PBS, washed in PBS and processed for FISH. FISH was performed as described in (Zuleger et al. 2013). Briefly, cells were permeabilized, cleared of RNA by RNase A and dehydrated with an ethanol series. DNA was denatured and captured in this state by a second ice-cold ethanol dehydration series. Coverslips were then annealed overnight to labelled BAC or whole chromosome probes. After washing probes were visualized with Alexa Fluor-conjugated Streptavidin/anti-digoxigenin antibodies and total DNA visualized with DAPI. To identify the nuclear lamina, coverslips were stained with a lamin B2 antibody before FISH. Coverslips were mounted in Vectashield (Vector Labs), images of nuclear midplanes acquired distance between genes and the nuclear lamins, or genes and other loci, were determined using Imaris.

### Microarrays

RNA from Resting and Activated Jurkat T-cells was extracted with TRI-reagent (Invitrogen). After quality testing using an Agilent Bioanalyzer (RNA 6000 Nano total RNA kit, Agilent) the RNA was converted to biotin labelled-cRNA using the TotalPrep RNA Amplification Kit (Ambion, AMIL1791) following the manufacturer's instructions. The quality of cRNA was then confirmed again using the Agilent Bioanalyzer. Quality analysis and hybridization of the cRNA was performed at the Wellcome Trust Clinical Research Facility in Edinburgh. For each experiment, three biological replicates were hybridized to Illumina whole genome gene expression arrays (HumanHT-12 v4 BeadChip). Hybridizations were carried out using an Illumina Beadstation. The subsequent microarray data were quantile normalized using the free statistics package R (R Development Core Team 2010) using the Bioconductor package Limma (Smyth 2005). Differentially expressed transcripts were selected with a log2 ratio above 0.5 in absolute value using moderated F-statistics adjusted for a false discovery rate of 5% (Benjamini and Hochberg 1995).

### DamID

DamID was performed as in (Vogel et al. 2007). Briefly, Resting Jurkat T-cells were transduced with Dam-Lamin B1‐ or Dam only-encoding lentiviruses over 24 h in the presence of 10ug/ul protamine sulphate. Transduced cells were then incubated with Raji B-cells ± SEE for a further 48 h after which genomic DNA was extracted and processed into libraries for next generation sequencing (Beijing Genomics). Sequenced reads were mapped to human Hg19 genome and the log2(Lamin B1/Dam) value determined for all genomic *DpnI* fragments in resting and activated cells, obtaining 2.2 and 2.1-fold genome coverage respectively. LADs were determined using a peak finder function in the BioConductor package DNAcopy, based in a circular binary segmentation algorithm (Seshan VE and Olshen A (2016). *DNAcopy: DNA copy number data analysis*. R package version 1.46.0). IP and PI regions were then identified as regions displaying differential LAD association.

### Hi-C Data Processing

The processed contact frequency maps are taken directly from the Rao et al. 2014 paper and displayed using the Juicebok Hi-C viewer (Rao et al. 2014). The “TADs” we refer to are in fact Contact Domains determined by Rao et al. 2014 and extracted from the publically available bed file. Where TADs tracks are shown in Figure 5, we have used ES cell TAD data from Dixon et al, 2012 and stated this in the text and figure legend. Additionally, the long-range interaction between the putative enhancer and CD200/BTLA that is displayed on Figure 7 was identified by Rao et al. using their HICCUP algorithm. This algorithm identifies loops between two points of DNA by determining the relative signal intensity between the interaction coordinate and its surrounding pixels. As such, interactions are registered as significant relative to other nearby interaction coordinates, which are subject to a similar distance separation. Finally, to determine the overlap between contact domains or A/B compartments with the Jurkat LADs we extracted the domain coordinates from bed files created in Rao et al. 2014. In this paper both compartments and contact domains were determined from normalized contact matrices adjust for the distance dependency decay of contact frequency.

## Supplemental information

Plasmids, antibodies, lentiviruses, bioinformatics, list of FISH probes and more detailed DamID and FISH procedures are described in Supplemental Methods.

8 Supplemental Figures, and Supplemental Methods.

## Data access

DamID and gene expression datasets are available at GEO (Accession Numbers to be added). For this work RNA-seq and H3K4me1 and H3K27ac ChIP-seq data for primary CD4^+^ T-cells was downloaded from the Epigenetics Roadmap Consortium (Bernstein et al. 2010; Kundaje et al.2015) and GM12878 cell RNA-seq from the Encode Consortium. Coordinates of H3K27ac-defined putative enhancer regions were extracted from Hnisz et al, 2013 while the positions of GM12878 Contact Domains and Sub-Compartments were extracted from Rao et al, 2014.

## Acknowledgements

We would like to thank Dr Peter Meinke and Ruth Shah for critical reading of the manuscript, Lior Pytowski and Anastasia Sokolidi for assistance with taking images, Shelagh Boyle for assistance with probe labelling, Dr David Kelly for microscopy support, and Louise Evenden at the Wellcome Trust Clinical Research Facility for microarray analysis. MIR was supported by a Wellcome Trust PhD Studentship and funding for this work was provided by Wellcome Trust grants 095209 to ECS and 092076 for the Centre for Cell Biology.

## Author contributions

Conceived of study and designed experiments, MIR and ECS; performed experiments, MIR, RC, AS; analyzed data, MIR, JdlH and ARWK; wrote the paper, MIR and ECS.

---

Supplemental Figures

Supplemental Methods

Supplemental References

## Supplemental Figures

**Figure S1.**
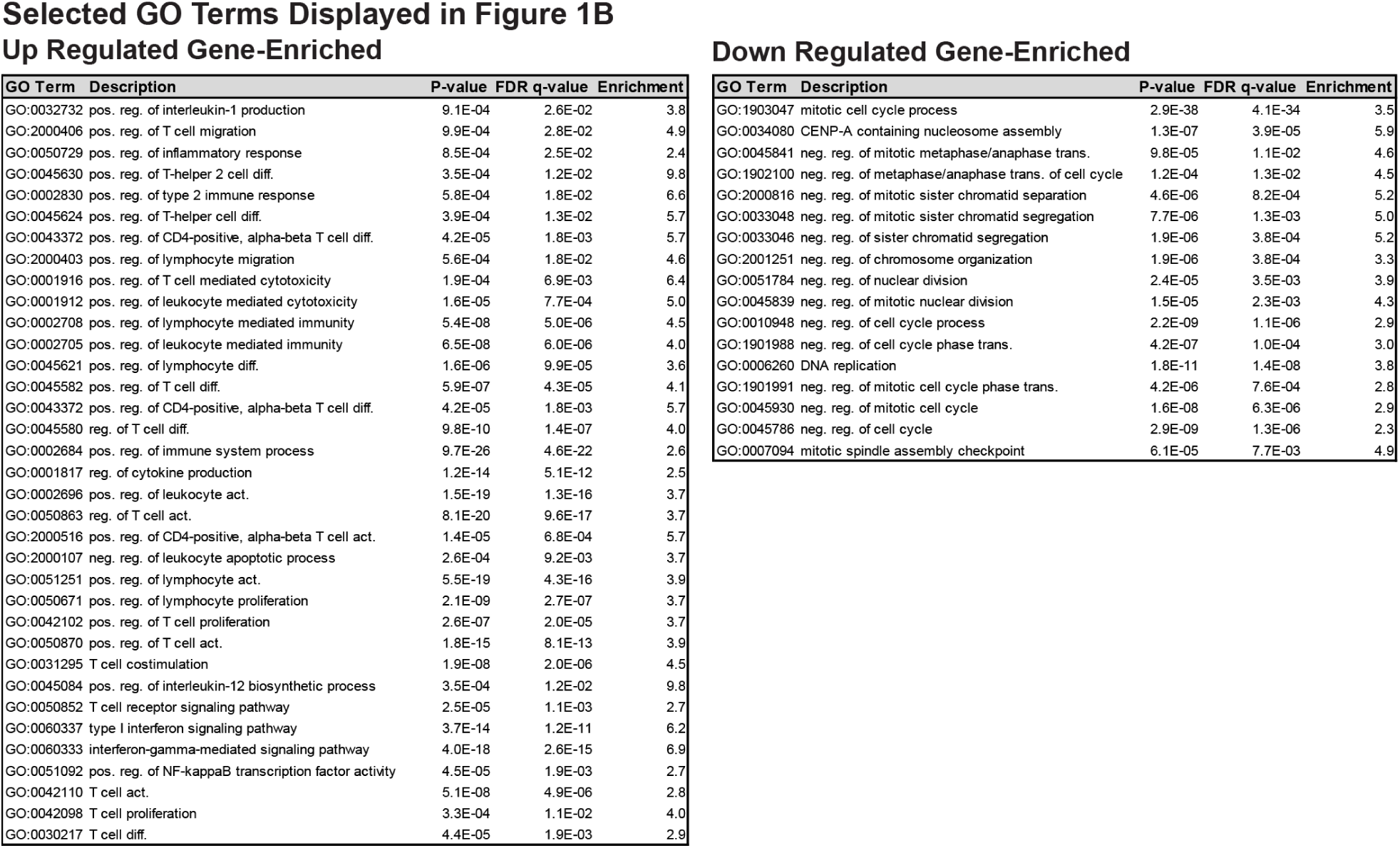
Summary of enriched GO terms categories used for Figure 1. See also Fig. 1.

**Figure S2.**
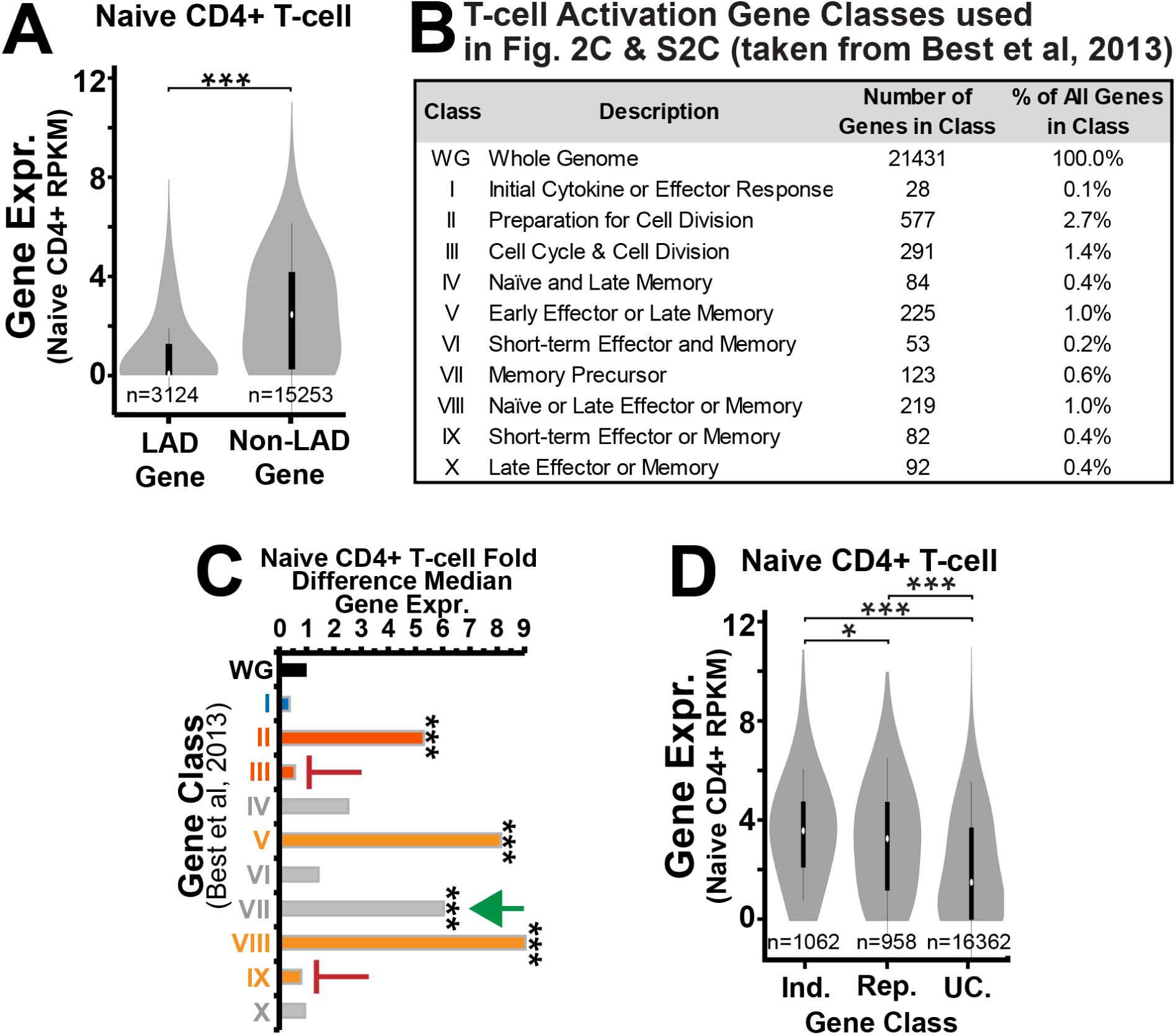
Lamina association is related to gene expression and function in naïve CD4^+^ T-cells. (*A*) Violin plot showing the expression of genes in naïve CD4^+^ T-cells that were in LAD or non-LAD regions in the resting Jurkat T-cells. Expression is in reads per Kb of transcript per Million mapped reads (RPKM) based on RNA-Seq data from naïve CD4^+^ T-cells. Genes in LADs in resting Jurkat cells are also less expressed than non-LAD genes in naïve CD4^+^ T-cells. (*B*) List of extracted T-cell activation gene classes identified by Best et al, 2013 through their transcriptional dynamics during an extensive T-cell activation time course. (*C*) Fold differences in gene expression in naïve CD4^+^ T-cells relative to the whole genome (WG) for the 10 categories of genes published by Best et al. Like resting Jurkat T-cells, classes II, V, and VIII are transcriptionally active in naïve CD4^+^ T-cells, and thus likely depleted from LADs also in the primary cells. By contrast, classes III and IX do not display elevated expression while class VII does. (*D*) Violin plot of the naïve CD4^+^ T-cells expression levels of genes that were induced (Ind.) or repressed (Rep.) during Jurkat T-cell activation. The distributions are similar to those for the Jurkat data shown in Figure 2. For A, B and E, naïve CD4^+^ T-cell RNA-seq data was extracted from the Epigenomics Roadmap Consortium Project (Kundaje et al. 2015). For A, C and D, significance of difference in gene expression was determined by Dunn test for multiple comparison testing after a Kruskal Wallis significant test. \**P* < 0.05, \*\**P* < 0.01 and \*\*\**P* < 0.001. See also Fig. 2.

**Figure S3.**
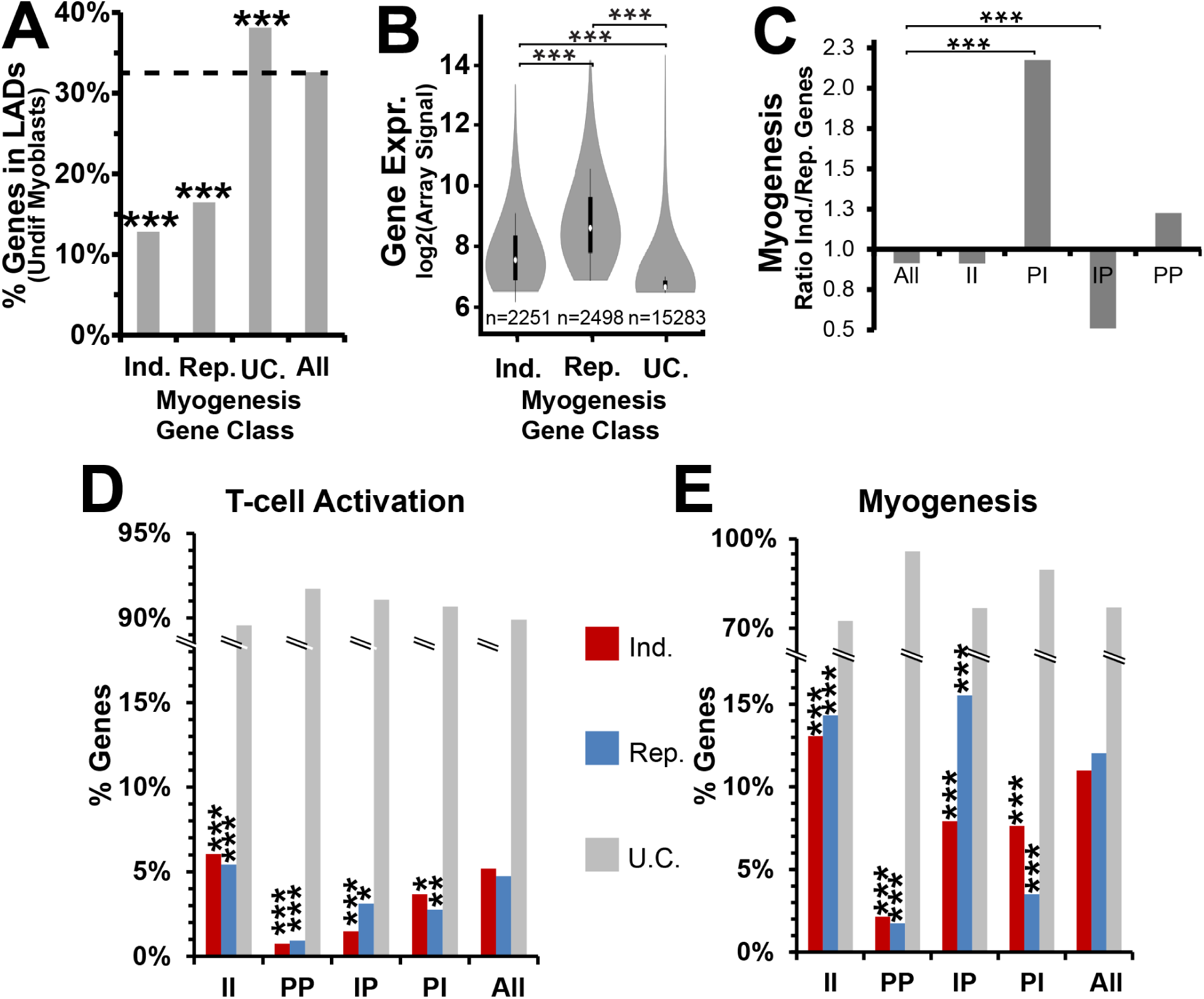
Comparison of T-cell activation‐ and myogenesis-induced gene expression and repositioning changes. (*A*) Bar plot similar to that in Figure 2D showing the showing the percentage of genes occurring in LADs in myoblasts for different expression categories: genes induced (Ind.) or repressed (Rep.) at least 1.4 fold during C2C12 myogenesis, unchanged genes (UC.) and all genes in the undifferentiated myoblasts. (*B*) Violin plot of the C2C12 myoblast expression level of genes induced, repressed, or unchanged during C2C12 myogenesis. (*C*) Bar plot demonstrating ratio of genes induced (Ind.) and repressed (Rep.) during C2C12 myogenesis for the whole genome (All), PI, IP, II and PP regions. (*D, E*). Bar graphs displaying gene expression change behaviour (Ind, Rep and UC) of IP, PI, II, PP and all genes during T-cell activation (*D*) and myogenesis (*E*). For both T-cell activation and myogenesis more PI genes are activated and repressed while more IP genes are repressed than activated. However, fewer repositioning genes display gene expression changes during T-cell activation than in myogenesis. All myogenesis DamID and gene expression data was re-analyzed from Robson et al, 2016. For A, C, D and E statistical significance was determined by Fisher tests. For B, significance of differences in gene expression was determined by Dunn test for multiple comparison testing after a Kruskal Wallis significant test. \**P* < 0.05, \*\**P* < 0.01 and \*\*\**P* < 0.001. See also Fig. 2.

**Figure S4.**
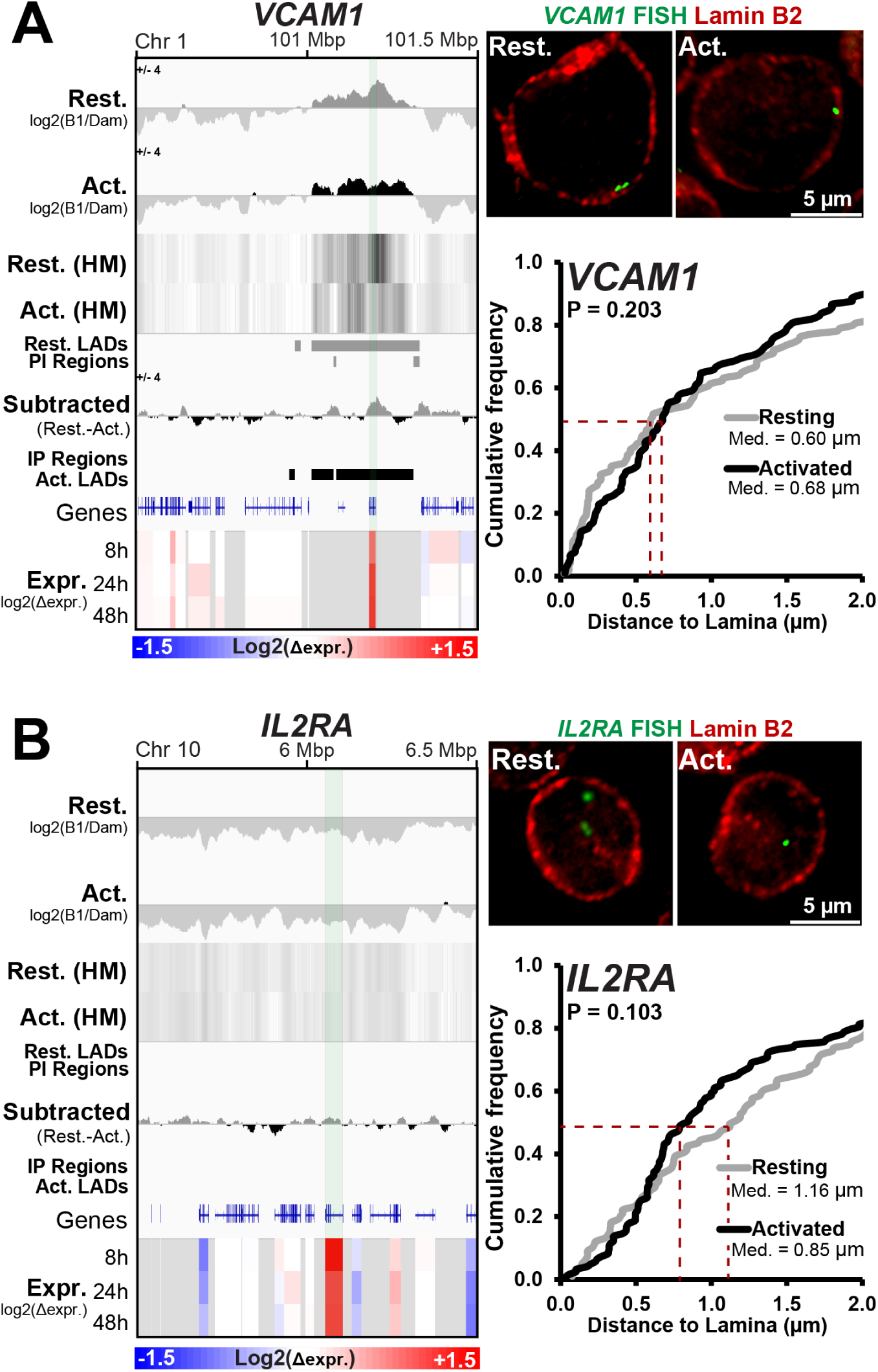
Genes maintaining positioning in LADs or non-LADs in both resting and activated Jurkat T-cells. (*A and B*) Genome browser views for the genomic region surrounding *VCAM1* and *IL2RA*, respectively, and quantification of loci positioning relative to lamin B2. DamID signal intensities, identified LADs, IP and PI regions and microarray gene expression changes for Resting and Activated Jurkat T-cells are shown. (*A*) The lamina-associated *VCAM1* locus, which contains no IP and PI regions, displays no significant release from the periphery during T-cell activation. (*B*) The consistently non-LAD *IL2RA* locus is significantly further from the periphery and displays no significantly altered positioning following activation. For quantification statistics, the position of loci in the activated sample was compared to the resting sample by KS tests. \**P* < 0.05, \*\**P* < 0.01 and \*\*\**P* < 0.001. See also Figs. 2 and 3.

**Figure S5.**
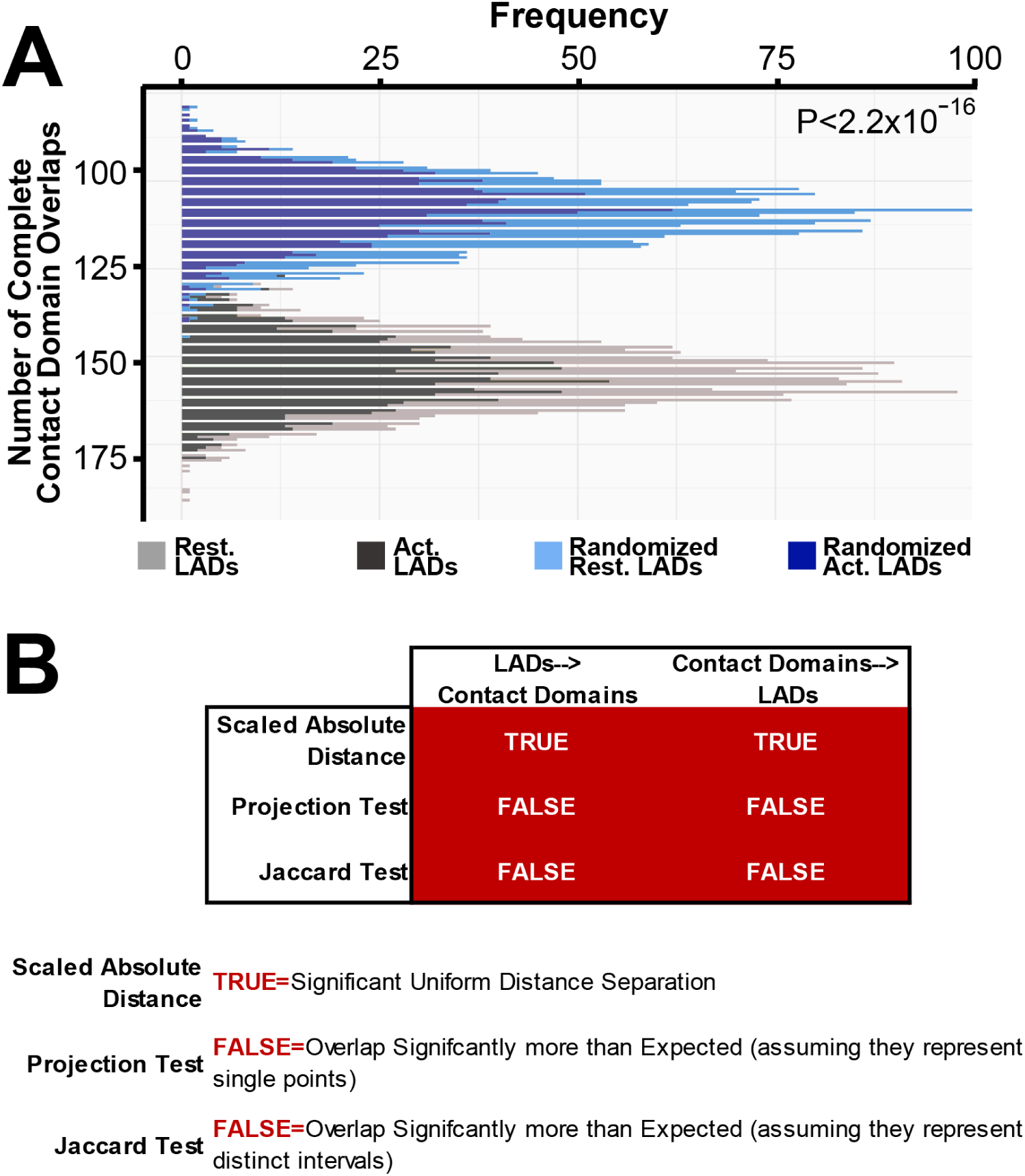
LADs are frequently encompassed within Contact Domains. (*A*) Histogram displaying the observed frequency of Jurkat LADs being completely encompassed within GM12878 Contact Domains compared to that expected in a randomly shuffled genome. 500 LADs were randomly selected and the frequency of complete envelopment within a Contact Domain determined. The positions of the 500 selected LADs were then randomly shuffled and frequency of complete envelopment re-determined. This process was then repeated 1,000 times, allowing all LADs of the genome to be examined multiple times, and the results plotted. LADs are completely enveloped by Contact Domains significantly more frequently than expected. Statistical significance for both resting and activated LADs was determined by comparing the distribution observed to that found in the randomly LAD-shuffled control by KS test. (*B*) Results of three test outputs from GenometriCorr analysis examining the relationship between GM12878 Contact Domains and resting Jurkat T-cell LADs. Note analysis is two directional with a query each time being contrasted to a reference. Thus for each comparison two results generated; LADs‐‐>Contact Domains and Contact Domains‐‐>LADs. For both directions results that reject the null hypothesis and support significant correlation or overlap are highlighted in red and explained in the legend within the figure. See also Fig. 5.

**Figure S6.**
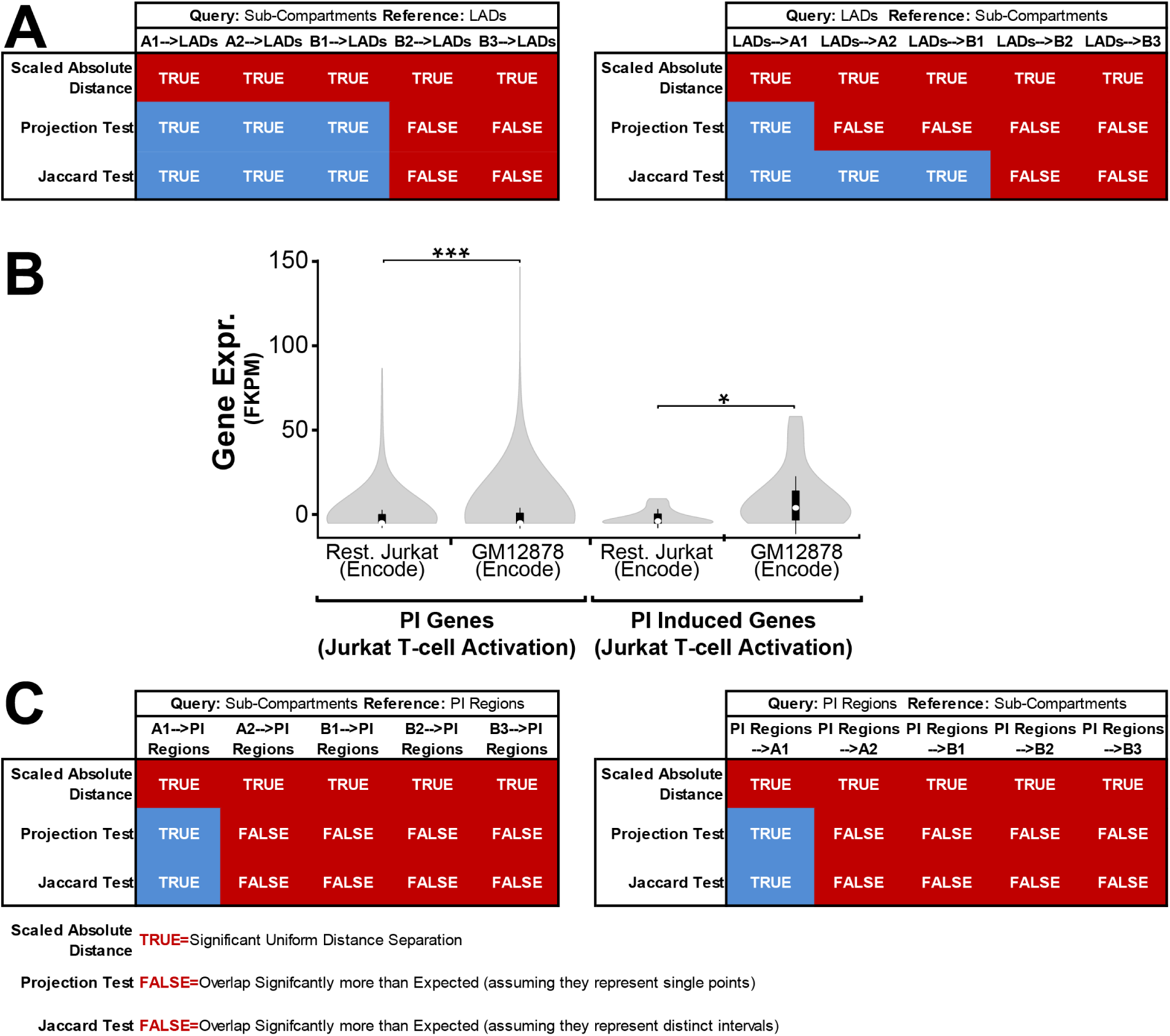
Genes induced and released from LADs during Jurkat T-cell activation also display elevated expression in GM12878. (*A*) GenometriCorr analysis of resting Jurkat LADs with GM12878 sub-compartments. Note that each test contrasts a query feature to a reference feature so for each pair of features, two results are obtained. LADs and all sub-compartments displayed distinct uniform distance separations (Scale Absolute Distance), regardless of the directionality of the test. Similarly by the Jaccard test, B2 and B3 Sub-compartments and LADs also display greater overlap than expected by chance, regardless of direction (B2 and B3‐‐>LADs and LADs‐‐>B2 and B3; Jaccard Tests). However, in the Projection tests, LADs also additionally displayed greater than expected overlap with the A2 and B1 subcompartments (LADs‐‐>A2 and B1) but not vice versa (A2 and B1‐‐>LADs). Hence the relationship between sub-compartments and LADs may be asymmetrical. (*B*) Violin plots showing the comparison of RNA-seq expression in ENCODE resting Jurkat and GM12878 cells for genes induced and released from the periphery in PI regions during Jurkat T-cell activation. Genes in PI genes, as well as PI genes induced during T-cell activation, both display elevated expression in GM12878 cells. For quantification, the expression of genes in ENCODE Resting Jurkat was compared to that in ENCODE GM12878 cells by KS tests. \**P* < 0.05, \*\**P* < 0.01 and \*\*\**P* < 0.001. (*C*). GenometriCorr analysis of PI regions with GM12878 sub-compartments. PI regions displayed greater than expected overlap with the A2, B1, B2 and B3 subcompartments, regardless of test directionality.

**Figure S7.**
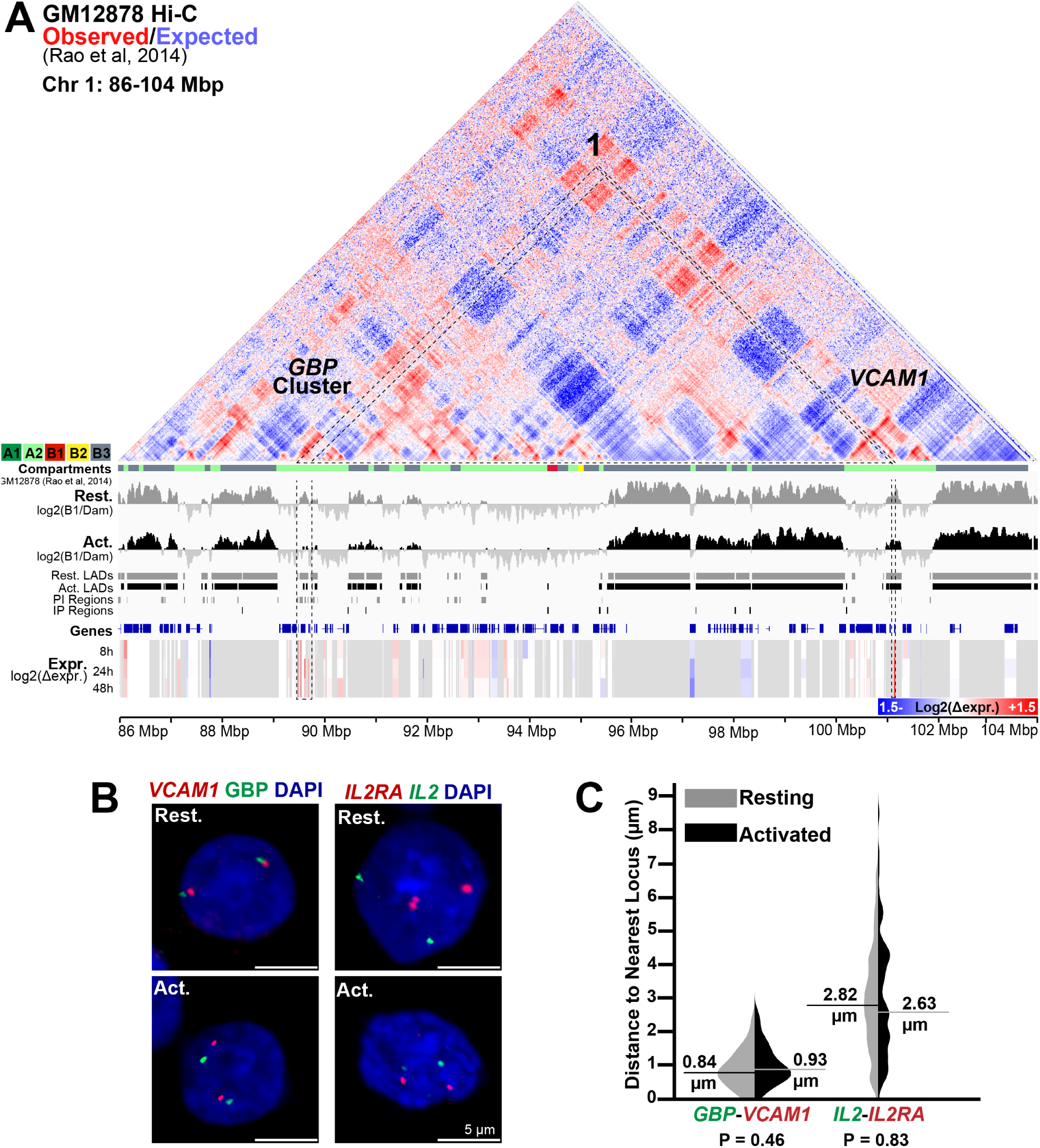
Loci that do not both release from LADs remain physically separated In activated Jurkat T-cells. (*A*) Genome browser view of a 17 Mb region of chromosome 1 with GM12878 cell observed/expected Hi-C interactions and sub-compartments (Rao et al. 2014) and DamID signal intensities, identified LADs, IP and PI regions and microarray gene expression changes for Resting and Activated Jurkat T-cells displayed as in Figure 7. The GBP gene cluster and *VCAM1* display no observable Hi-C interaction. (*B and C*) Representative images and quantification respectively of inter-loci distances in resting and activated T-cells for the *VCAM1* and the *GBP* gene cluster, located 11 Mb apart, and the *IL2RA* and *IL2* loci, located on separate chromosomes. The distance between *IL2RA* and *IL2*, or the more spatial proximal *VCAM1* and *GBP* gene cluster, remains unchanged during T-cell activation. For quantification statistics, the positions of loci in the activated sample were compared to the resting sample by KS tests. \**P* < 0.05, \*\**P* < 0.01 and \*\*\**P* < 0.001. See also Fig. 7.

**Figure S8.**
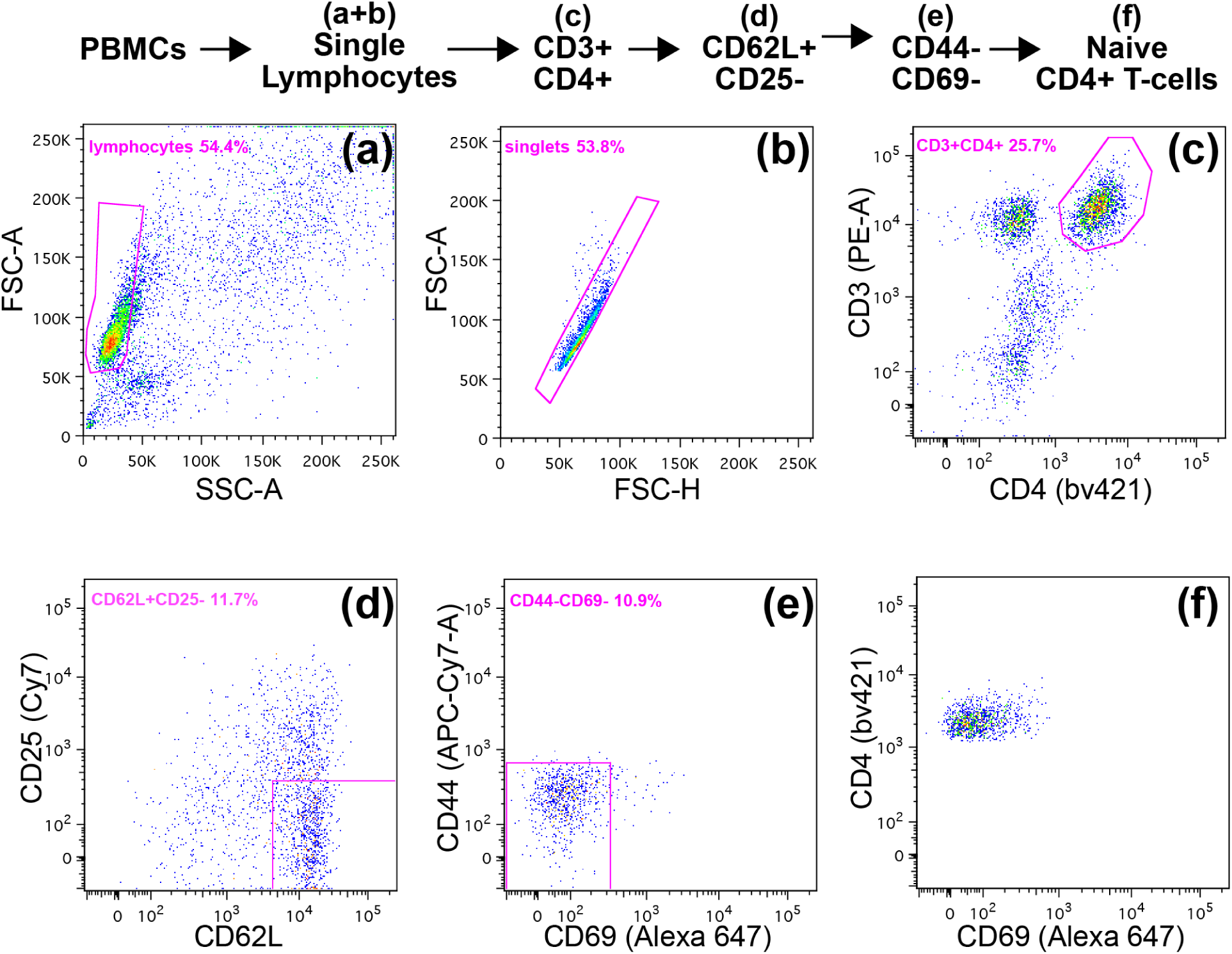
FACs gating scheme for isolation of naïve CD4^+^ T-cells. Single lymphocytes were first sorted from PBMCs, necrotic cells and debris using forward light scatter area (FSC-A), forward light scatter height (FSC-H) and side light scatter area (SSC-A) gates (*A*) and (*B*). CD4^+^ T-cells were then selected by CD3 and CD4 staining (*C*). Naïve cells with a lack of activation markers were isolated by sequential gates for CD25 negative and CD62L positive cells (*D*) followed by CD44 negative and CD69 negative cells (*E*). The last panel (*F*) shows the final CD4 positive and CD69 negative population. Hence, a population of pure CD3^+^, CD4^+^, CD25^-^, CD62L+, CD44^-^ and CD69^-^ naïve T-cells were isolated. See also Fig. 8.

## Supplemental methods

### Plasmid and cell sources

psPAX2 and pMD2.G were a gift from Justina Cholewa-Waclaw (Adrian Bird, WTCCB, Edinburgh). pLgw Dam-V5-Lamin B1 and pLgw V5-Dam were a gift from the van Steensel laboratory. Jurkat T-lymphocyte clone E6-1 was purchased from ATCC while Raji lymphoblastoid cells were a gift from Professor Vicente Andrés (Centro Nacional de Investigaciones Cardiovasculares. Madrid, Spain).

### Antibodies

Rabbit anti-Lamin B2 (Schirmer et al. 2001) antibody was used at a 1:400, dilution. For visualization of primary antibodies for immunofluorescence, donkey anti-rabbit pig secondary antibodies conjugated to a variety of Alexa Fluor^®^ dyes were used (Molecular Probes, Invitrogen). For the detection of biotin‐ or digoxigenin-labelled probes in FISH experiments, streptavidin (Molecular Probes, Invitrogen) or anti-digoxigenin antibodies (Jackson labs) conjugated to Alexa Fluor^®^ dyes were used, respectively.

### Lentivirus generation and transduction

Lentiviruses encoding DamID constructs were generated as described in Robson et al, 2016 with several modifications. Briefly, non-replicative lentiviruses were generated by transfection of ∼6 million 293FT cells plated in an 8.5 cm diameter tissue culture plate with 2.8 μg pMD2.G, 4.6 μg psPAX2 and 7.5 μg of the construct-specific transfer vector using 36 μl lipofectamine 2000 in 3 ml Opti-MEM as per the manufacturer’s instructions. After 16 h 293FT media was replaced. 48 h later the virus containing supernatant was aspirated, cleared of cellular debris by centrifugation for 10 min at 3,500 rpm and followed by filtration through a 0.45 μm^2^ low protein binding PES syringe filter (Millipore, SLHP003RS). Viruses were then concentrated by ultracentrifugation at 55,000 × *g* for 75 min at 4^o^C in a JA-25.5 rotor and resuspended in an appropriate volume of Opti-MEM. If not used immediately, aliquots were frozen at -80°C. Transduction was performed in the presence of 10 μg/ml protoamine sulphate.

## Bioinformatics

### Jurkat T-cell Activation Gene Expression Analysis

Genes differentially expressed in resting and activated Jurkat T-cells analyzed for Biological Process and Cellular Compartment GO-term enrichment using Gene Ontology enrichment analysis and visualization tool GOrilla (Salmon and Trono 2007). Genes upregulated during T-cell activation were significantly enrichment in terms positively associated with T-cell activation while those downregulated were enriched in terms associated with negative regulation of the cell cycle. A full list of GO-terms enriched in genes upregulated and downregulated during T-cell activation can be found in Supplementary Figure S1. To demonstrate the gene expression changes associated with GO-categories, specific GO-terms that either support T-cell activation or negatively regulate cell cycle were selected and displayed in Fig. 1. These terms are shown in Fig. S1.

### DamID (more detailed)

DamID was performed as described in (Vogel et al. 2007). Briefly for each DamID sample two million Jurkat cells were transduced with ^1^/_10_ of a Dam methylase encoding lentiviral preparation in the presence of 10 μg/ml protamine sulphate in a 3.5 cm diameter tissue culture dish. After 24 h cells were pelleted at 200 × *g* for 4 min and resuspended in fresh suspension cell growth media. Dam-control or Dam-Lamin B1-transduced Jurkat T-cells were then incubated with untransduced Raji B cells ±SEE and left a further 48 h, diluting cells in appropriate volumes of suspension cell growth media to sustain logarithmic growth, before proceeding with DNA extraction.

DamID sample processing was then performed as described in Vogel et al. Briefly, DNA was extracted from cells using the DNeasy tissue lysis kit (Qiagen) as per manufacturer’s instructions. 2.5 μg of extracted DNA was then digested by *DpnI* (NEB) and, following heat inactivation of *DpnI*, was ligated to the DamID adaptor duplex (dsAdR) generated from the oligonucleotides AdRt (5’-CTAATACGACTCACATAGGGCAGCGTGGTCGCGGCCGA-GGA-3’) and AdRb (5’-TCCTCGGCCG-3’) after which DNA was further digested by *DpnII*. To amplify DNA sequences methylated by the Dam methylase, 5 μl of *DpnII* digested material was then subjected to PCR in the supplied buffer in the presence of the 1.25 μM Adr-PCR primer (5’-GGTCGCGGCCGAGGATC-3’), 0.2 mM dNTPs and 1X of the Advantage cDNA polymerase (Clontech, cat. no. 639105). PCR was performed as described in Table 1.

**Table 1.**
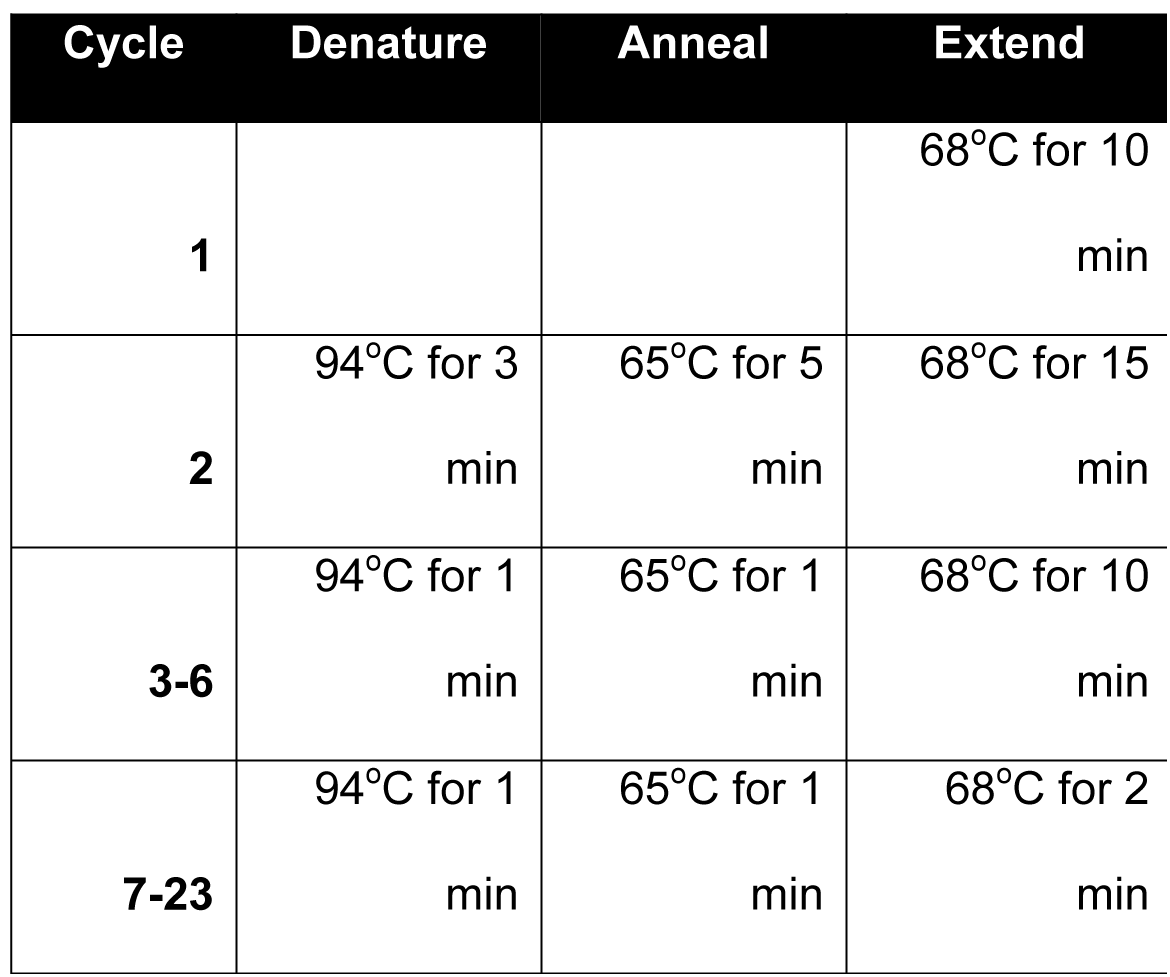
PCR program for DNA amplification in DamID.

| Cycle | Denature | Anneal | Extend |
| --- | --- | --- | --- |
| **1** |  |  | 68°C for 10 min |
| **2** | 94°C for 3 min | 65°C for 5 min | 68°C for 15 min |
| **3-6** | 94°C for 1 min | 65°C for 1 min | 68°C for 10 min |
| **7-23** | 94°C for 1 min | 65°C for 1 min | 68°C for 2 min |

Following PCR, the distribution of amplified DNA fragments was checked on agarose gels (Fig. 1). If samples were of sufficient quality DNA was purified on QIAquick PCR purification columns (Qiagen) and then concentrated to the required concentration by precipitation. To generate the 2 μg of material required of next generation sequencing it was found an average of 6-8 PCR reactions were required per sample.

### DamID Sequencing and Analysis

DamID sample libraries were prepared for next generation sequencing by fragmentation followed by ligation to sequencing adaptors. Libraries were then sequenced by 90 bp paired end (90PE) sequencing in reactions with 5 samples per well. DamID sequences were aligned to the human Hg19 genome using the Burrows-Wheeler Aligner software bwa-mem (Quinlan and Hall 2010). Subsequent processing was performed using R (R Core Team (2015). R: A language and environment for statistical computing. R Foundation for Statistical

Computing, Vienna, Austria. URL https://www.R-project.org/) and Bedtools (Eden et al. 2009). Resting and Activated Jurkat DamID data was quantified at a DpnI fragment level by counting the number of reads that overlapped each DpnI-flanked (GATC) genomic fragment for each pair of Dam-alone and Dam-LaminB1 samples. The log2 ratios between Dam-LaminB1 and Dam-alone were calculated for each DpnI fragment, and the resulting values quantile normalized in R using the BioConductor Limma package (Li and Durbin 2009) to allow a more quantifiable sample comparison.

### Lamina Associated Domains (LADs) Definition

Normalized data were smoothed by a ∼ 15Kb window in order to stabilize noise, while maintaining the same number of data points. To identify LADs we used a circular binary segmentation algorithm in the Bioconductor package DNAcopy (Seshan VE and Olshen A (2016). *DNAcopy: DNA copy number data analysis*. R package version 1.46.0) using the default parameters. The positive signal tracts were extracted and merged if they were <8Kb apart. We removed anything below 15Kb from our LAD list. This threshold was chosen empirically after careful examination of the LAD traces as most of the smaller LADs are too close to noise. We counted a gene as being in a LAD if it overlapped with one.

### Identification of Genomic Regions with Altered Peripheral Association

To identify genomic regions with differential frequencies of association to the periphery, we subtracted each set of LADs from each other to obtain a set of regions uniquely present in each set. We filtered this set to eliminate anything smaller than 8Kb or with a signal difference (log2) lower than 0.3. This filter significantly reduces the number of false positives due to slight differences in the definition of LAD ends, or due to regions with very low intensity.

### Analysis of LAD and/or PI region intersect with GM12878 Contact Domains and Compartments

To accurately compare the frequency with which LADs are completely enveloped within Contact Domains, 500 LADs were randomly selected and the number of times a LAD was found completely within a Contact Domain determined. We then randomized the location of each LAD per chromosome in the down-sampled files and re-counted the frequency of envelopment. This process was repeated 1000 times to ensure all LADs in the genome were analyzed multiple times. Both the scripts and the data used for figure S3 can be retrieved from the github site: https://github.com/AlastairKerr/Robson2016). The significance of the difference in distribution between observed and randomized was determined by KS test.

In order to determine their relative enrichment in GM12878 sub-compartments, the degree of overlap of each individual LADs or PI regions was determined and contrasted to that expected in a genome where LADs or PI regions had been randomly shuffled. The significance of the distribution between observed and randomized was determined by KS test.

### Fluorescence In Situ Hybridization (more detailed)

For FISH experiments Resting and Activated Jurkat cells were pelleted at 200 × g for 5 min and resuspended in PBS. Cells were then plated onto poly-lysine coated coverslips, left for 5 minutes and then fixated in 4% para-formaldehyde, 1X PBS for 10 min at room temperature. After aging coverslips for several days, cells were permeabilized for 6 min with 0.2% Triton-X-100 in PBS, followed by 3 washes in PBS. If antibody staining was required coverslips were blocked with 1% BSA prior to sequential incubations with primary and secondary antibodies. After washing, antibodies were fixed for 45 s in 2% paraformaldehyde, PBS. Cells were next pre-equilibrated in 2X SCC and treated with RNase A (100μg/ml) at 37°C for 1 h. Following washing in 2X SCC, cells were dehydrated with a 70%, 85% and 100% ethanol series. Coverslips were then air dried, heated to 70°C and submerged into 85°C preheated 70% formamide, 2X SSC (pH 7.0) for 21 min. A second ethanol dehydration series was then performed using -20°C 70% ethanol for the first step. Coverslips were air dried and 150-300 ng biotin-/digoxigenin-/fluorophore-labelled probe was added in hybridization buffer (50% formamide, 2X SSC, 1% Tween20, 10% Dextran Sulphate) containing 6 μg human Cot1 DNA (Invitrogen) and sheared salmon sperm DNA and incubated at 37°C for 24 h in a humidified chamber. After incubation, the coverslips were washed four times for 3 min each in 2X SSC at 50°C followed by four times for 3 min each in 0.1X SSC at 65°C. Coverslips were then pre-equilibrated in 2X SSC, 0.1% Tween-20 and blocked with 4% BSA before incubating for 1 h at room temperature with Alexa Fluor^®^conjugated-Steptavidin/anti-dioxigenin antibodies and 4,6-diamidino-2 phenylindole, dihydrochloride (DAPI) at 2 μg/ml. Coverslips were subsequently washed 3 times in 2X SSC, 0.1% Tween-20 at 37°C and mounted on slides in Vectashield (Vector Labs). BACs were ordered as indicated in Table 2.

**Table 2.** List of FISH probes.

| **Locus** | **Clone ID** | **Chr** | **Start Coordinate** | **End Coordinate** |
| --- | --- | --- | --- | --- |
| GBP i | WI2-1164H23 | 1 | 89,225,254 | 89,263,381 |
| GBP ii | WI2-1912I12 | 1 | 89,441,903 | 89,486,300 |
| GBP iii | WI2-2752H5 | 1 | 89,657,420 | 89,702,009 |
| GBP iv | WI2-526L3 | 1 | 89,876,247 | 89,917,750 |
| GBP v | WI2-1217K14 | 1 | 90,091,207 | 90,135,261 |
| VCAM1 i | WI2-2673A12 | 1 | 100,736,026 | 100,781,301 |
| VCAM1 ii | WI2-1322N21 | 1 | 100,934,306 | 100,971,590 |
| VCAM1 iii | WI2-1574P13 | 1 | 101,119,857 | 101,157,755 |
| VCAM1 iv | WI2-569L23 | 1 | 101,303,334 | 101,342,416 |
| VCAM1 v | WI2-481B14 | 1 | 101,490,965 | 101,526,327 |
| Putative<br>Enhancer | WI2-898A6 | 3 | 98,250,119 | 98,286,966 |
| NFKBIZ | WI2-645D14 | 3 | 101,546,513 | 101,590,433 |
| CBLB | WI2-628H6 | 3 | 105,507,827 | 105,545,557 |
| BTLA | WI2-1965N10 | 3 | 112,175,307 | 112,212,883 |
| CD200 | WI2-526H19 | 3 | 112,039,557 | 112,083,246 |
| IL2 | CH17-337H11 | 4 | 123,272,797 | 123,467,914 |
| TNFA | CH17-119I1 | 6 | 31,527,402 | 31,735,928 |
| IL2RA | CH17-215L21 | 10 | 5,959,657 | 6,190,768 |
| CCND2 | WI2-2236P14 | 12 | 4,261,002 | 4,303,424 |

### Microscopy

Images were acquired on a Nikon TE-200 microscope using a 1.45 NA 100x objective, Sedat quad filter set, PIFOC Z-axis focus drive (Physik Instrubments) and a CoolSnapHQ High Speed Monochrome CCD camera (Photometrics) run by Metamorph image acquisition software.

